# Inhibitory control and error processing in Obsessive-Compulsive Disorder: A mega-analysis of task-based fMRI data by the ENIGMA-OCD consortium

**DOI:** 10.1101/2025.10.22.683868

**Authors:** Nadža Džinalija, Odile A. van den Heuvel, H. Blair Simpson, Iliyan Ivanov, Ana Araújo, Srinivas Balachander, Jan Beucke, Daniel Brandeis, Silvia Brem, Willem Bruin, Jan Buitelaar, Miguel Castelo-Branco, Sunah Choi, Goi Khia Eng, Sophie M.D.D. Fitzsimmons, Lydia Fortea, Miquel A. Fullana, Rosa Grützmann, Bjarne Hansen, Chaim Huijser, Niels T. de Joode, Norbert Kathmann, Christian Kaufmann, Minah Kim, Kathrin Koch, Jun Soo Kwon, Jie Xin Lim, Ignacio Martinez-Zalacain, Jose M. Menchon, Laurens A. van de Mortel, Janardhanan C. Narayanaswamy, Olga Therese Ousdal, Tjardo S. Postma, Daniela Rodriguez-Manrique, Daan van Rooij, Venkataram Shivakumar, Carles Soriano-Mas, Emily R. Stern, Sophia I. Thomopoulos, Anders L. Thorsen, Enric Vilajosana, Susanne Walitza, Lea Waller, Ysbrand D. van der Werf, Guido van Wingen, Stella J. de Wit, Dan J. Stein, Paul M. Thompson, Chris Vriend, Ilya M. Veer

## Abstract

**Objective:** Obsessive-compulsive disorder (OCD) is a chronic condition in which impaired inhibitory control and excessive error monitoring may contribute to the maintenance of obsessions and compulsions. This mega-analysis investigates neural activation during response inhibition and error processing using adult and pediatric data from the ENIGMA-OCD consortium and the ABCD study.

**Methods:** Individual participant data was uniformly processed using HALFpipe to extract statistical maps for response inhibition and error processing contrasts. Bayesian multilevel models were used to assess regional and whole-brain effects of OCD, with additional analyses examining the association between the OCD clinical profile and task-related activation.

**Results:** Across inhibitory control tasks, both individuals with OCD and control participants showed robust activation in regions implicated in response inhibition and error processing. During response inhibition, compared to controls, adults with OCD showed stronger somatomotor cortex activation, while children with OCD showed stronger occipital cortex activation. Children with likely OCD from the ABCD cohort showed reduced activity in the frontoparietal network in the anterior insula/frontal operculum region. During error processing, relative to controls, adults with OCD showed weaker activation in fronto-striatal regions, while children with OCD showed stronger activation in frontoparietal and attention networks. Greater OCD symptom severity was associated with weaker task-related activation in adults and stronger activation in children during response inhibition.

**Conclusion:** Case-control differences in brain activation during inhibitory control varied by age group and task contrast. Symptom severity emerged as the main clinical correlate of activation during inhibition, suggesting that inhibitory control deficits in OCD may be both state-dependent and developmentally specific.

## Introduction

A hallmark of obsessive-compulsive disorder (OCD) is the inability to inhibit thoughts (obsessions) and behaviors (compulsions). The ability to withhold mental and physical behaviors, known as response inhibition(1), is a sub-construct of the *cognitive systems* domain of the Research Domain Criteria (RDoC(2)) and is impaired in OCD. Individuals with OCD show deficits in inhibitory control tasks such as the Stop-signal and Flanker tasks, which engage the ventral cognitive circuit, including the dorso-lateral prefrontal cortex (dlPFC), the pre-supplementary motor area (pre-SMA), and the dorsal caudate(3). Meta-analytic evidence points to weaker activation in OCD in the anterior cingulate cortex (ACC), thalamus, caudate, supramarginal gyrus (SMG), anterior insula and frontal operculum (aI/fO), orbitofrontal cortex (OFC) and occipital lobe during successful inhibition. Stronger activation was found during failed inhibition in the ACC, pre-SMA, aI/fO, and PFC, implying hyper-vigilance to error(4). Clinical characteristics further modulate these processes, as antidepressants have been linked to decreased activation during response inhibition(5), while weaker activation has been associated with greater symptom severity(6). Early onset of OCD was found to correlate with worse executive control performance(7), though its relationship with neural activation during inhibition is unclear. Characterizing the neural basis of inhibitory control in OCD is difficult because prior meta-analyses often conflated inhibitory control with broader executive functions(8–11) and lacked the individual-level data to examine clinical heterogeneity.

The ENIGMA-OCD consortium (Enhancing Neuro-Imaging and Genetics through Meta-Analysis) conducts mega-analyses by pooling participant-level neuroimaging data, offering greater statistical power to detect case-control differences and explore clinical heterogeneity than meta-analyses that aggregate results at the study level(12, 13). We previously investigated domain-relevant activation in negative valence(14) and executive function(15) in the first group-level analyses combining participant-level whole-brain fMRI data from different task paradigms. By capturing individual-level effects, this mega-analysis approach provides a more nuanced view of OCD-related brain alterations than traditional coordinate-based meta-analyses.

Here we investigated inhibitory control (response inhibition and error monitoring) in OCD as a distinct domain, disentangling it from higher-order executive functions such as working memory or planning. We pooled individual participant data from 17 adult and pediatric samples globally, all of whom completed inhibitory control tasks during MRI. We further investigated clinical characteristics of OCD, such as medication usage, symptom severity, and age of onset. We hypothesized that individuals with OCD would show weaker activation than controls in brain regions supporting inhibitory control during response inhibition, but stronger activation in error-monitoring regions during errors. We expected that weaker response inhibition-related activation would be associated with a medicated status, greater symptom severity, and an early onset of OCD.

## Methods

### Study population

Sixteen samples (6 unpublished) across 10 countries were obtained from the ENIGMA-OCD consortium (Table S1), a global network with legacy neuroimaging data from OCD individuals and healthy controls (HCs). An additional sample came from the ABCD study, a population cohort of 9-11 year old US adolescents, from which we selected those with suspected OCD and age- and sex-matched HCs. OCD diagnosis in ENIGMA-OCD samples was determined by DSM criteria using diagnostic tools administered by trained personnel, while severity was measured using the (Children’s) Yale-Brown Obsessive-Compulsive Scale ((C)Y-BOCS;(16, 17)). OCD is assessed in the ABCD cohort by parent-report on the Kiddie-Schedule for Affective Disorders (K-SADS;(18, 19)). Because OCD diagnosis was not verified by a clinician in the ABCD sample, this sample is considered at high likelihood or at-risk for OCD. HCs were free of psychopathology and were not taking psychotropic medication. Participants gave informed consent and protocols were approved by local review boards, permitting re-use of de-identified data by ENIGMA-OCD.

### Inhibitory control tasks

Participants from each sample completed an fMRI inhibitory control task assessing different aspects of inhibition: three samples used the Flanker task (measuring interference control), two used action restraint tasks (one Go/No-go, one eye-blink suppression task), and twelve used the stop-signal task (measuring action cancellation; Table 1). Interference control involves inhibiting intrusion of distracting or irrelevant stimuli, action restraint requires suppressing a prepotent or instinctual response, and action cancellation inhibits an already initiated motor response. Despite these differences, each task requires inhibitory control of an automated response, allowing us to define one common inhibition versus response contrast, as well as an error processing contrast where possible (15 tasks).

**Table 1.**
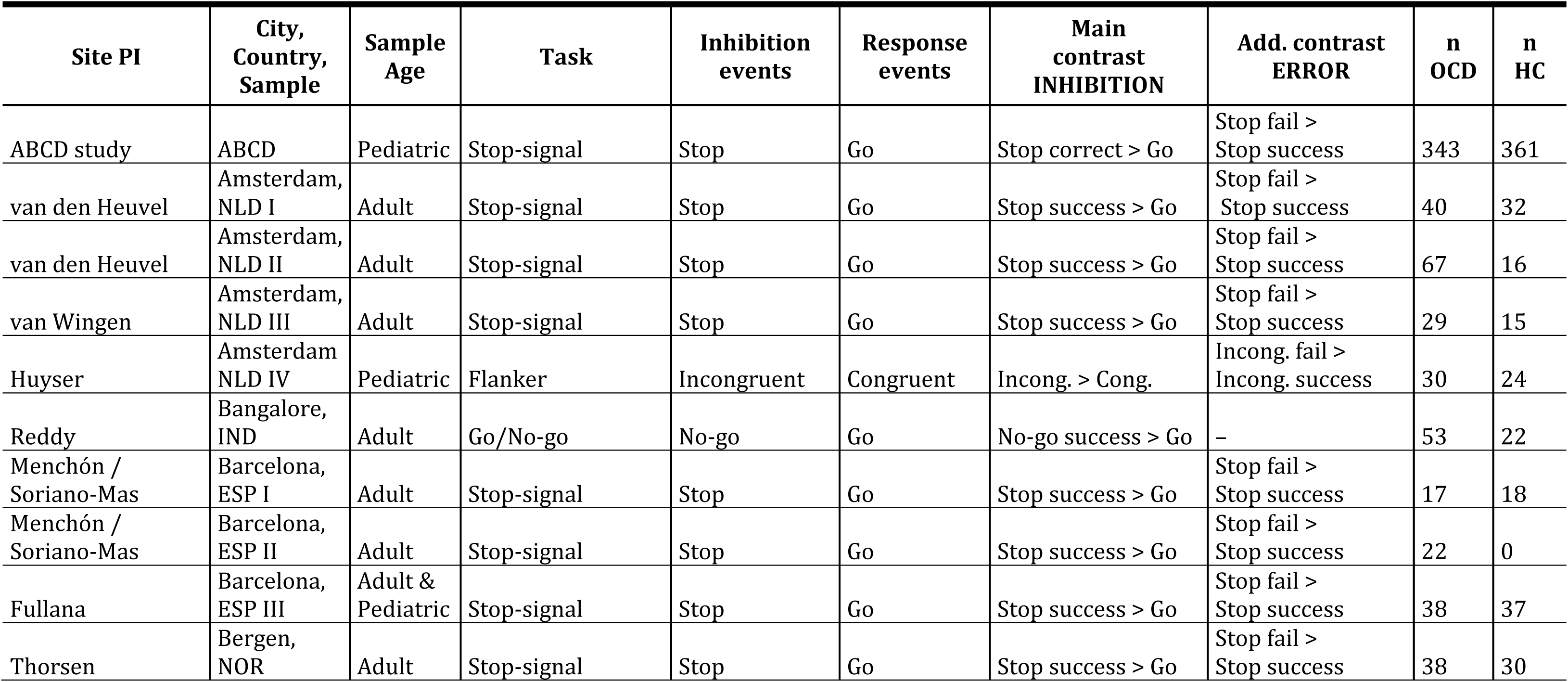
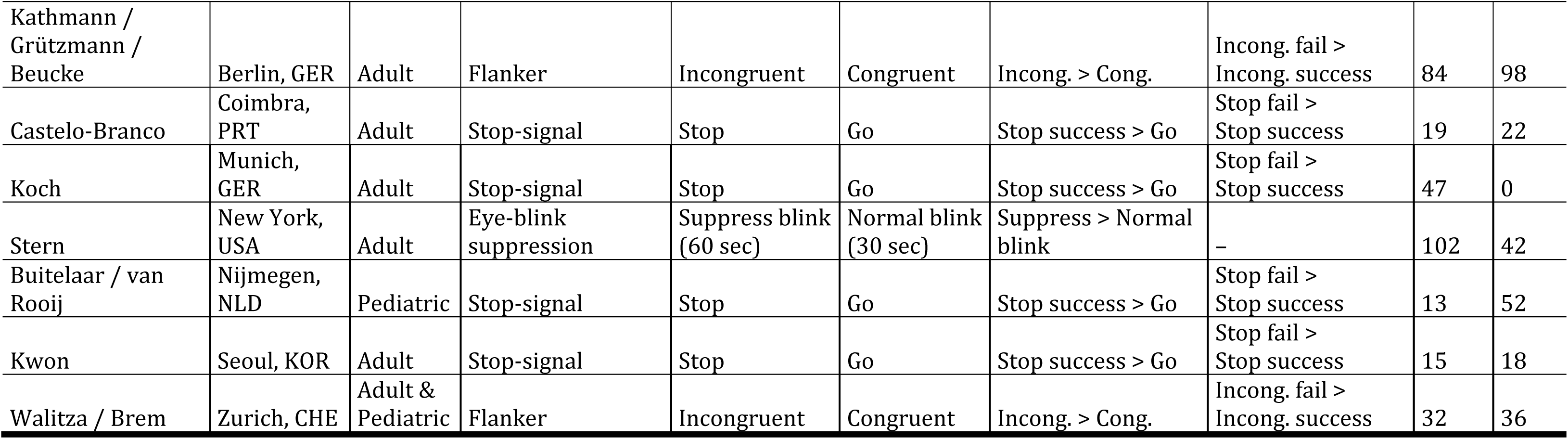
Tasks included in mega-analysis. All tasks modeled events using event-related regressors with zero duration (i.e., stick functions) except the eye-blink suppression task which used a block design.

| Site PI | City,<br>Country,<br>Sample | Sample<br>Age | Task | Inhibition<br>events | Response<br>events | Main<br>contrast<br>INHIBITION | Add. contrast<br>ERROR | n<br>OCD | n<br>HC |
| --- | --- | --- | --- | --- | --- | --- | --- | --- | --- |
| ABCD study | ABCD | Pediatric | Stop-signal | Stop | Go | Stop correct > Go | Stop fail ><br>Stop success | 343 | 361 |
| van den Heuvel | Amsterdam,<br>NLD I | Adult | Stop-signal | Stop | Go | Stop success > Go | Stop fail ><br>Stop success | 40 | 32 |
| van den Heuvel | Amsterdam,<br>NLD II | Adult | Stop-signal | Stop | Go | Stop success > Go | Stop fail ><br>Stop success | 67 | 16 |
| van Wingen | Amsterdam,<br>NLD III | Adult | Stop-signal | Stop | Go | Stop success > Go | Stop fail ><br>Stop success | 29 | 15 |
| Huyser | Amsterdam<br>NLD IV | Pediatric | Flanker | Incongruent | Congruent | Incong. > Cong. | Incong. fail ><br>Incong. success | 30 | 24 |
| Reddy | Bangalore,<br>IND | Adult | Go/No-go | No-go | Go | No-go success > Go | – | 53 | 22 |
| Menchón /<br>Soriano-Mas | Barcelona,<br>ESP I | Adult | Stop-signal | Stop | Go | Stop success > Go | Stop fail ><br>Stop success | 17 | 18 |
| Menchón /<br>Soriano-Mas | Barcelona,<br>ESP II | Adult | Stop-signal | Stop | Go | Stop success > Go | Stop fail ><br>Stop success | 22 | 0 |
| Fullana | Barcelona,<br>ESP III | Adult &<br>Pediatric | Stop-signal | Stop | Go | Stop success > Go | Stop fail ><br>Stop success | 38 | 37 |
| Thorsen | Bergen,<br>NOR | Adult | Stop-signal | Stop | Go | Stop success > Go | Stop fail ><br>Stop success | 38 | 30 |
| Kathmann /<br>Grützmann /<br>Beucke | Berlin, GER | Adult | Flanker | Incongruent | Congruent | Incong. > Cong. | Incong. fail ><br>Incong. success | 84 | 98 |
| Castelo-Branco | Coimbra,<br>PRT | Adult | Stop-signal | Stop | Go | Stop success > Go | Stop fail ><br>Stop success | 19 | 22 |
| Koch | Munich,<br>GER | Adult | Stop-signal | Stop | Go | Stop success > Go | Stop fail ><br>Stop success | 47 | 0 |
| Stern | New York,<br>USA | Adult | Eye-blink<br>suppression | Suppress blink<br>(60 sec) | Normal blink<br>(30 sec) | Suppress > Normal<br>blink | – | 102 | 42 |
| Buitelaar / van<br>Rooij | Nijmegen,<br>NLD | Pediatric | Stop-signal | Stop | Go | Stop success > Go | Stop fail ><br>Stop success | 13 | 52 |
| Kwon | Seoul, KOR | Adult | Stop-signal | Stop | Go | Stop success > Go | Stop fail ><br>Stop success | 15 | 18 |
| Walitza / Brem | Zurich, CHE | Adult &<br>Pediatric | Flanker | Incongruent | Congruent | Incong. > Cong. | Incong. fail ><br>Incong. success | 32 | 36 |

### MRI Image acquisition and processing

Sites contributed previously-acquired legacy data (Table S1). Image data processing was harmonized across samples (see Supplemental Methods), including preprocessing the structural and functional images and extracting the task contrasts of interest using the open-source containerized Harmonized AnaLysis of Functional MRI pipeline (HALFpipe version 1.2.2;(20, 21)). Using default settings within HALFpipe, pre-processing for structural images included skull stripping and spatial normalization and for functional images included motion correction, slice time correction (if slice acquisition order was known), susceptibility distortion correction (if fieldmaps were available), coregistration, spatial normalization, denoising with six (rigid-body) motion parameters, and smoothing with a 6mm FWHM Gaussian kernel.

We excluded participants with mean framewise displacement >1.0 mm. Processed data was quality controlled at each site according to harmonized guidelines (see Supplemental Methods).

#### First-level contrast parameter estimate maps

The first-level contrast of interest across all tasks compared inhibition/suppression conditions (*stop* trials in the Stop-signal, Go/No-go and eye-blink suppression tasks, and *incongruent* trials in the Flanker tasks) with simple response conditions (*go* trials in the Stop-signal, Go/No-go, and eye-blink suppression tasks, and *congruent* trials in the Flanker tasks) (Table 1). The error contrast modeled errors of commission on *stop* or *incongruent* trial, contrasted with successful inhibition.

## Analyses

The analysis and hypotheses were pre-registered at osf.io/mhq8t and analysis scripts are available at github.com/nadza-dz/task-based-fMRI-processing-pipeline-ENIGMA-OCD.git. We used the same processing and analysis pipeline as Dzinalija et al. on negative valence(14) and executive function(15).

### ROI analyses

We investigated regions of interest (ROIs) identified in Norman et al.’s(4) meta-analysis of inhibitory control in HC samples. For response inhibition, these regions comprised the aI/fO, SMA, superior parietal lobe (SPL), superior occipital lobe, premotor cortex (PMC), and dlPFC; for error processing ROIs were the aI/fO, ACC, occipital lobe, and SMG (Table S2). Peak activation coordinates from Norman et al. were transformed to MNI152 NLIN 2009c (asymmetric) space, and 5 mm spheres were created around each coordinate. Mean activation within each ROI was extracted from the z-statistic maps of each participant’s first-level inhibition and error contrasts. Multiple runs were aggregated at participant-level before extracting the contrast estimates.

Rather than applying separate linear models to each ROI, we analyzed all regions simultaneously using a Bayesian multilevel model (RBA, v1.0.10;(22)), which accounts for shared information across brain regions within a participant in one statistical model. Unlike null hypothesis significance testing, the Bayesian approach allows us to directly assess hypothesis credibility while accounting for complex dependencies in the data and mitigating multiple testing issues. For a more detailed discussion of this approach, refer to Dzinalija et al.(10) and Chen et al.(22). We used a noninformative Gaussian prior and four Markov chains (4,000 iterations each), with convergence defined as R̂ <1.1. The credibility of an effect was inferred from the area under the curve of the positive posterior distribution (see Supplemental Methods), summarized as the positive posterior probability (P+). We interpreted P+ values as moderate evidence of an effect (<0.10 or >0.90), strong evidence (<0.05 or >0.95), or very strong evidence (<0.025 or >0.975)(22).

All analyses were conducted in three separate groups: ENIGMA-OCD adults (age ≥18), ENIGMA-OCD children (age <18), and the ABCD sample (ages 9-10). We first assessed the main effect of inhibitory control across tasks and all individuals using an intercept model. Our primary effect of interest was the case-control effect. We also investigated clinical subgroups through three sets of analyses: 1) pairwise comparisons between early-onset OCD (age of onset <18), late-onset (age of onset ≥18), and HCs, restricting to adults within ENIGMA-OCD; 2) pairwise comparisons of medicated (any psychotropic medication) individuals, unmedicated individuals, and HCs in both ENIGMA-OCD and ABCD; and 3) associations between symptom severity (continuous (C)Y-BOCS) and activation in ENIGMA-OCD (not available in ABCD). Sample, age, and sex were included as covariates in all models. Because ABCD data were collected at 18 different sites, each site was treated as a separate sample to account for scanner effects.

### Sensitivity analyses

To assess the robustness of the response inhibition results, several sensitivity analyses were conducted. Sample effects were examined with a leave-one-sample-out analysis for each ROI analysis in the adult ENIGMA-OCD samples. We then assessed the influence of sample correction method by comparing our main case-control analyses (which modeled sample as a covariate of no interest) with ComBat harmonization(23). Because each sample administered only one inhibitory control task, controlling for sample also implicitly controlled for task differences; nonetheless, we explicitly tested task effects by replacing sample with task as a covariate. We additionally restricted analyses to the stop-signal task in adult ENIGMA-OCD samples to assess whether results were consistent when focusing on a single, well-represented paradigm. Finally, to evaluate robustness across statistical frameworks, we compared Bayesian results for the case-control analyses with a frequentist multilevel mixed-effects linear model that included sample and participant as random effects.

### Whole-brain analyses

To implement the Bayesian multilevel model across the entire brain, analyses were performed within functionally defined anatomical regions using the Schaefer-Yeo 7-network 200-parcel atlas(24) for cortical areas and the Melbourne subcortical atlas scale 2(25) for subcortical structures. Each participant’s first-level contrast z-statistic maps were parcellated, and activation values were incorporated into the same Bayesian multilevel group analyses described earlier, provided that at least 30% of voxels in the parcel contained signal (Table S3). Sample, sex, and age were included as covariates of no interest.

## Results

### Participants

Our final ENIGMA-OCD sample included 563 adults with OCD (54.9% female, mean age=32.4±10.7), 348 adult HCs (52.0% female, mean age=31.5±10.1), 83 children with OCD (41.0% female, mean age=13.3±2.7), and 114 pediatric HCs (44.3% female, mean age=12.5±2.3) (Table 2, S4). Our ABCD sample consisted of 343 children with likely OCD (43.4% female, mean age=9.99±0.6) and 361 HC children (49.6% female, mean age=9.97±0.6). For exclusions (n=85 OCD, n=60 HCs) and per-analysis sample sizes see Supplemental Methods.

**Table 2.** Demographics of samples included in mega-analysis. Data are expressed as mean and standard deviation (SD) unless otherwise noted. Medication status was measured at the time of scan. OCD = obsessive-compulsive disorder, HC = healthy control; (C)Y-BOCS = (Children’s) Yale-Brown Obsessive-Compulsive Scale.

|  | Adult<br>ENIGMA-OCD samples |  |  | Pediatric<br>ENIGMA-OCD samples |  |  | ABCD<br>sample |  |  |
| --- | --- | --- | --- | --- | --- | --- | --- | --- | --- |
|  | OCD (n=563) | HC (n=348) | Statistics | OCD (n=83) | HC (n=114) | Statistics | OCD (n=343) | HC (n=361) | Statistics |
| **Sex** |  |  |  |  |  |  |  |  |  |
| n female | 309 (54.88%) | 181 (52.01%) | $X^2_{(1)} = 0.6,$<br>$p = 0.44$ | 34 (40.96%) | 51 (44.74%) | $X^2_{(1)} = 0.15,$<br>$p = 0.7$ | 149 (43.44%) | 179 (49.58%) | $X^2_{(1)} = 2.43,$<br>$p = 0.12$ |
| n male | 254 (45.12%) | 167 (47.99%) |  | 49 (59.04%) | 63 (55.26%) |  | 194 (56.56%) | 182 (50.42%) |  |
| **Age** | 32.41 (10.74) | 31.46 (10.11) | $T_{(909)} = -1.32,$<br>$p = 0.19$ | 13.25 (2.7) | 12.56 (2.33) | $T_{(195)} = -1.91,$<br>$p = 0.06$ | 9.99 (0.6) | 9.97 (0.58) | $T_{(702)} = -0.58,$<br>$p = 0.56$ |
| **Age of onset** |  |  |  |  |  |  |  |  |  |
| n onset <18 | 252 (44.76%) | - |  | 83 (100%) | - |  | 343 (100%) | - |  |
| n onset ≥18 | 190 (33.75%) | - |  | - | - |  | - | - |  |
| n missing data | 121 (21.49%) | - |  | - | - |  | - | - |  |
| **Medication** |  |  |  |  |  |  |  |  |  |
| n medicated | 274 (48.67%) | - |  | 11 (13.25%) | - |  | 60 (17.49%) | - |  |
| n unmedicated | 287 (50.98%) | - |  | 71 (85.54%) | - |  | 283 (82.51%) | - |  |
| n missing data | 2 (0.36%) | - |  | 1 (1.2%) | - |  | - | - |  |
| **(C)Y-BOCS** | 23.69 (6.2) | - |  | 22.5 (6.81) | - |  | - | - |  |
| n missing data | 3 (0.53%) | - |  | 5 (6.02%) | - |  | - | - |  |

### Task effects: Response inhibition and error processing

Across all groups, all ROIs showed very strong evidence of stronger activation during response inhibition (all P+≥0.91; Figure 1A-C) except the left PMC (P+=0.66) in adults. The right PMC (P+=0.01) and left SPL (P+=0.03) showed evidence for weaker activation in the adult group. There was very strong evidence of stronger activation in all ROIs during error processing across all tasks and groups (P+>0.96; Figure 1D-F), except in the right SMG (P+=0.2) and left STG (P+0.47) in the ABCD group.

**Figure 1.**
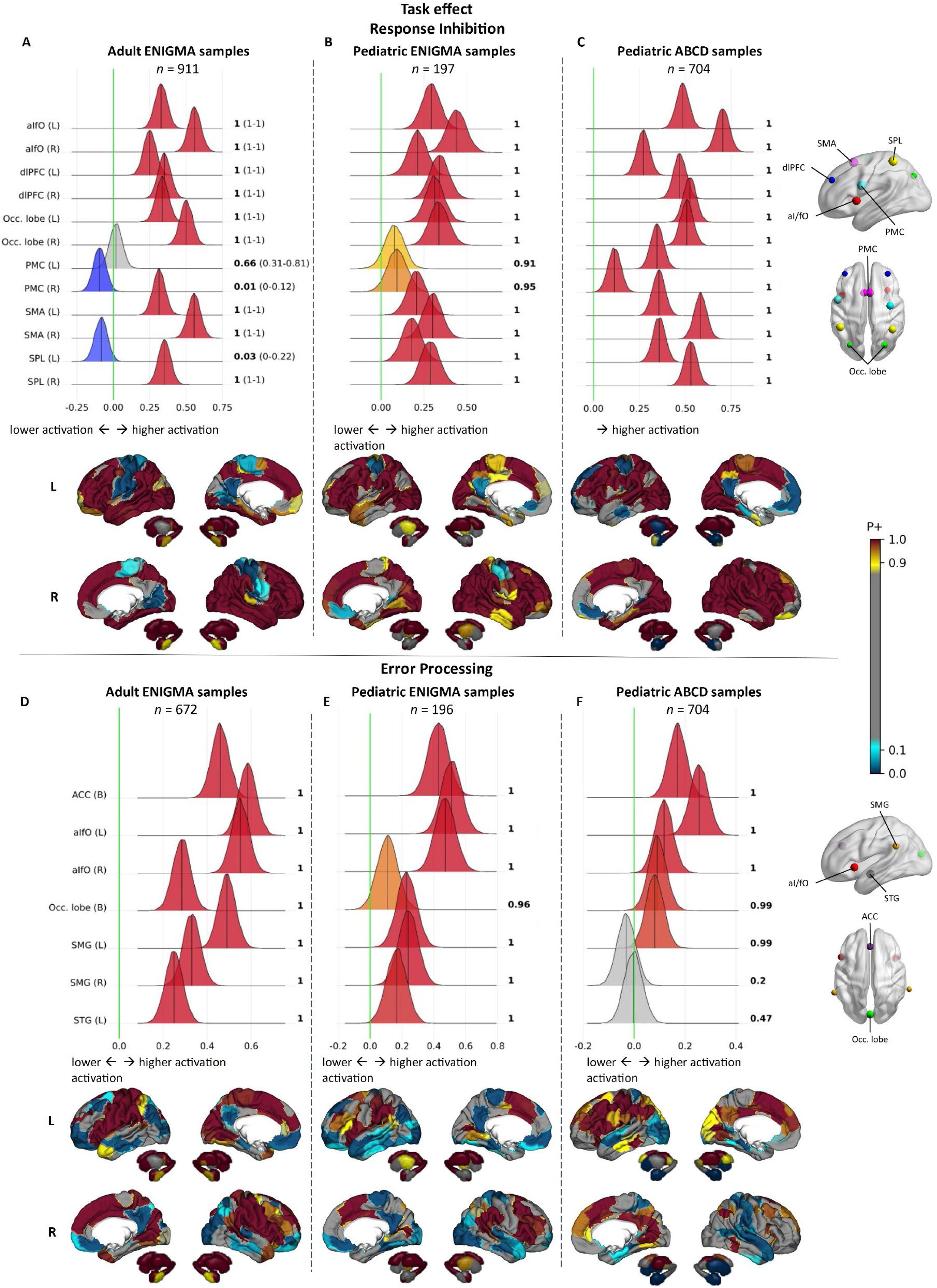
Main effect of response inhibition and error processing across all participants. Results from Bayesian multilevel analyses of group-level contrasts for response inhibition (upper figure) and error processing (lower figure). Analyses were performed across individuals with OCD and healthy controls combined, and across all tasks, separately within each set of samples: (A,D) Adult ENIGMA-OCD samples, (B,E) Pediatric ENIGMA-OCD samples, and (C,F) Pediatric ABCD samples. For each sample, the top panels display region-of-interest results across the predefined regions displayed to the right of the figure, while the bottom panel displays whole-brain results. In the ROI panels, posterior probability distributions express the credibility of an effect in each region. Next to each distribution the posterior probability of a positive effect (P+) is shown in bold, as well as the range of values this probability took on in leave-one-sample-out sensitivity analyses (where this analysis was carried out). Distributions to the right of the green no-effect line indicate stronger activation, while those to the left indicate weaker activation. Regions are color-coded to reflect the strength of evidence: (darker) red = stronger evidence for activation (P+ values > 0.90 indicate moderate to very strong evidence for a positive effect), (darker) blue = stronger evidence for deactivation (P+ values < 0.10 indicate moderate to very strong evidence for a negative effect), grey = no strong evidence of activation or deactivation. Values on the x-axis represent the difference in regional activation levels between experimental and baseline conditions for each contrast (expressed as difference in Z-scores). In the whole-brain panels, P+ values denote the probability of stronger brain activation in a given region of the Schaefer 200-parcel 7-network cortical atlas and Melbourne 32-region subcortical atlas. Displayed are lateral and medial views of the cortex and subcortex (amygdala, hippocampus, thalamus, nucleus accumbens, pallidum, putamen, and caudate). ACC = anterior cingulate cortex; aI/fO = anterior insula/frontal operculum; dlPFC = dorsolateral prefrontal cortex; L = left; Occ = occipital; PMC = primary motor cortex; R = right; SMA = supplementary motor area; SMG = supramarginal gyrus; SPL = superior parietal lobule; STG = superior temporal gyrus.

Whole-brain analyses showed a similar pattern across groups of stronger response inhibition-related activation of regions spanning the frontoparietal, dorsal and ventral attention, and visual networks, and weaker activity in somatomotor and default mode networks. During error processing, there was stronger activation in ventral attention regions, combined with weaker visual activation across groups, as well as stronger activation in somatomotor regions in adults.

### Inhibitory activation in OCD versus HCs

We did not find evidence for any activation differences between individuals with OCD and HCs in predefined ROIs during response inhibition in ENIGMA-OCD adults or children (Figure 2A-B). In the ABCD sample (Figure 2C), children with likely OCD showed moderate evidence for weaker activation than HCs in the right aI/fO (P+=0.07) and right SMA (P+=0.09). During error processing (Figure S2D), adults with OCD showed weaker activation than HCs, with strong evidence in the ACC (P+=0.03) and moderate evidence in the bilateral aI/fO and bilateral SMG (0.06<P+<0.08). No differences were found between OCD and HCs in the pediatric samples during error processing (Figure S2E-F).

**Figure 2.**
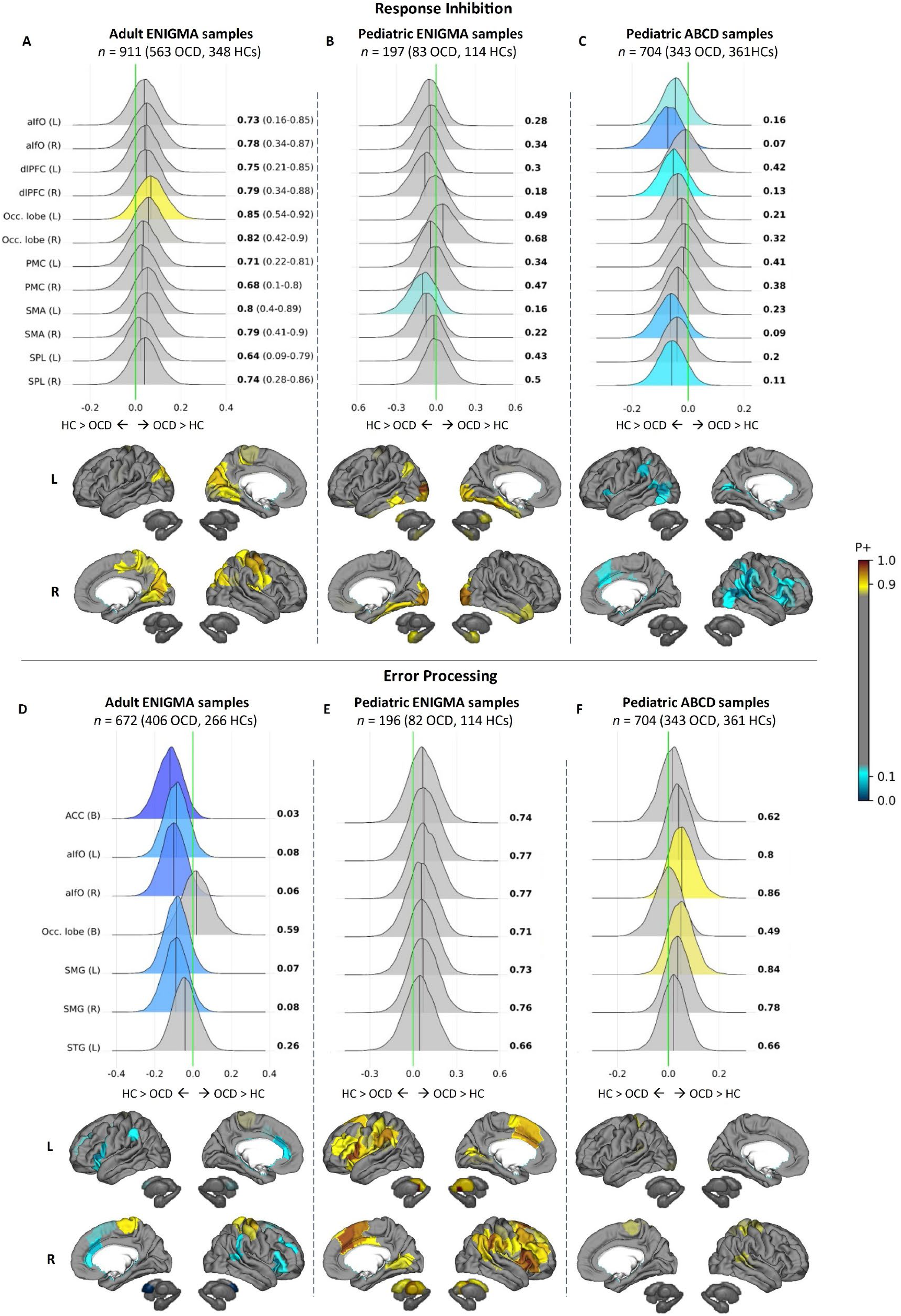
Case-control differences in response inhibition and error processing. Results from Bayesian multilevel analyses of response inhibition and error processing, contrasting individuals with OCD from healthy controls across each set of samples: (A,D) Adult ENIGMA-OCD samples, (B,E) Pediatric ENIGMA-OCD samples, and (C,F) Pediatric ABCD samples. For each sample, the top panels display region-of-interest results while the bottom panel displays whole-brain results. In the ROI panels, distributions to the right of the green no-effect line represent regions in which individuals with OCD show evidence for stronger activation than HCs. (Darker) red color represents regions in which individuals with OCD show moderate-to-very-strong evidence for stronger activation than HCs. Regions with posterior distributions to the left of the no-effect line show evidence for stronger activity in HCs than in OCD. (Darker) blue color represents regions in which HCs show moderate-to-very-strong evidence for stronger activation than OCD individuals. In gray-colored regions there is no evidence of a difference between HCs and OCD individuals. Values on the x-axis represent the difference in regional activation levels between HCs and OCD (expressed as difference in Z-scores). ACC = anterior cingulate cortex; aI/fO = anterior insula/frontal operculum; dlPFC = dorsolateral prefrontal cortex; L = left; Occ = occipital; PMC = primary motor cortex; R = right; SMA = supplementary motor area; SMG = supramarginal gyrus; SPL = superior parietal lobule; STG = superior temporal gyrus.

Whole-brain analyses in response inhibition revealed stronger activation in the somatomotor network in adults with OCD relative to HCs, while children with OCD in the ENIGMA-OCD sample showed stronger activation in select visual regions. During error processing, adults with OCD exhibited weaker activation in the anterior caudate and ventral attention regions relative to HCs. In contrast, ENIGMA-OCD children with OCD showed stronger activation in the thalamus, as well as regions of frontoparietal and attention networks relative to HCs.

### Clinical correlates of inhibitory control in OCD

#### Response inhibition

There was no evidence that age of OCD onset (early vs. late) or medication status (medicated vs. unmedicated) influenced brain activation during response inhibition in either children or adults in ENIGMA-OCD (Figures S1,S3). In contrast, in the ABCD cohort, medicated children with likely OCD showed moderate to strong evidence for stronger activation in the right dlPFC (P+=0.92) and bilateral SPL (0.91<P+<0.95) (Figure 3C). In adults with OCD (Figure 3A), moderate evidence linked higher symptom severity to weaker activation during response inhibition in the bilateral SPL, bilateral PMC, and left SMA (0.08<P+<0.10). This was accompanied by widespread weaker activation across nearly all whole-brain regions. In contrast, ENIGMA-OCD children showed an opposite pattern (Figure 3B), where higher symptom severity was associated with stronger activation in the bilateral SPL, bilateral occipital lobe, bilateral aI/fO, and bilateral dlPFC (P+>0.90). Whole-brain analyses confirmed this association, revealing widespread stronger activation with increasing symptom severity across frontoparietal, dorsal/ventral attention, visual, and nearly all subcortical regions.

**Figure 3.**
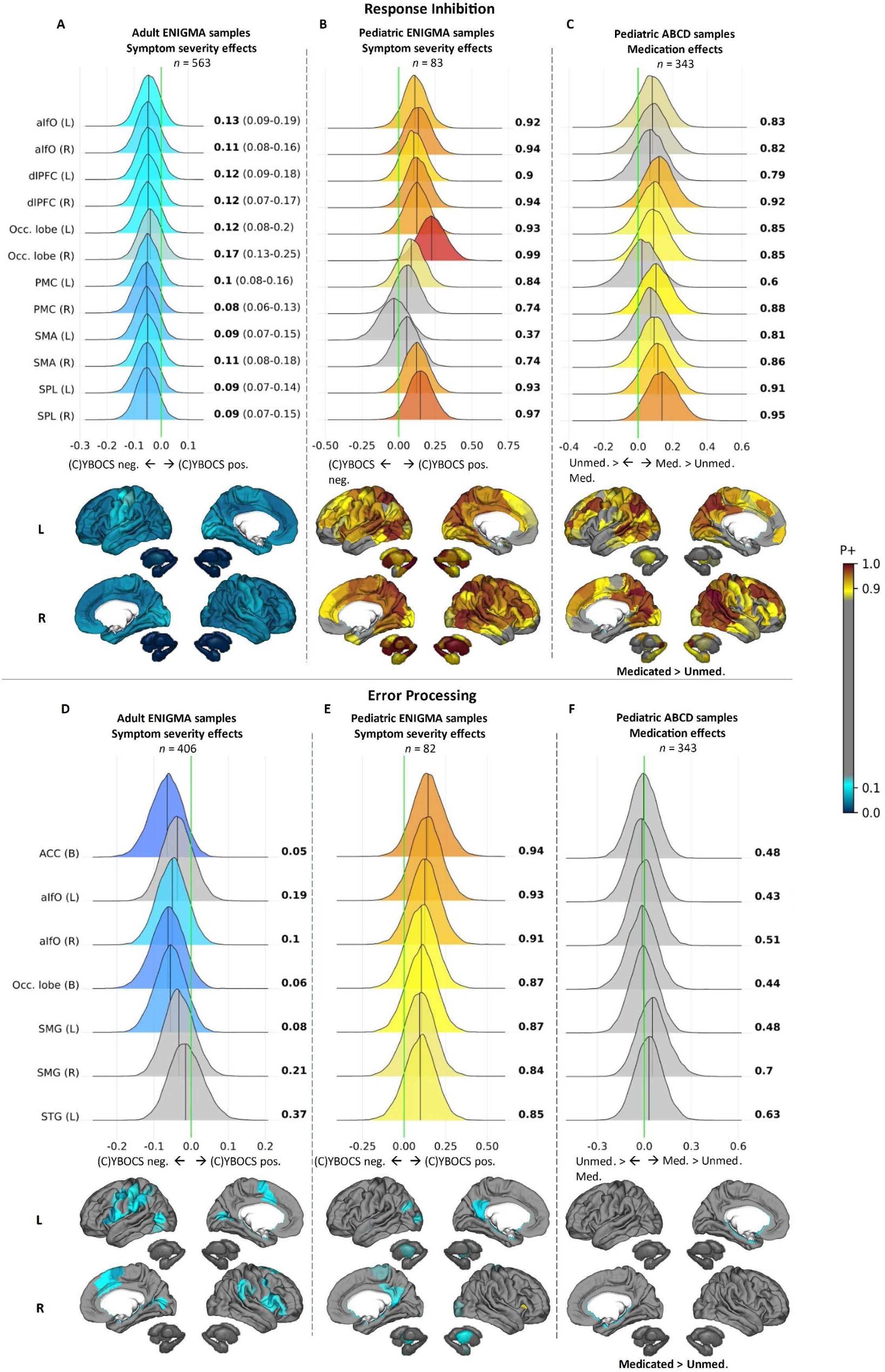
Effects of clinical features of OCD on response inhibition and error processing. Select region-of-interest and whole-brain effects of symptom severity and medication status during response inhibition and error processing, across each set of samples: (A,D) Adult ENIGMA-OCD samples, (B,E) Pediatric ENIGMA-OCD samples, and (C,F) Pediatric ABCD samples. Full reporting of all investigated effects can be found in Figures S1-S5. ACC = anterior cingulate cortex; aI/fO = anterior insula/frontal operculum; (C)YBOCS = (Children’s) Yale-Brown Obsessive-Compulsive Scale; dlPFC = dorsolateral prefrontal cortex; L = left; Occ = occipital; PMC = primary motor cortex; R = right; SMA = supplementary motor area; SMG = supramarginal gyrus; SPL = superior parietal lobule; STG = superior temporal gyrus.

#### Error processing

Findings for error processing in the predefined ROIs were broadly similar to response inhibition in adults with OCD (Figure S2), with no credible evidence of differences by age of onset or medication status. However, there was moderate evidence that higher symptom severity was linked to weaker activation in the ACC (P+=0.05), occipital lobe (P+=0.06) and left SMG (P+=0.08) (Figure 3C). Among children with OCD or likely OCD, medication status was not associated with activation differences during error processing in either the ABCD or ENIGMA-OCD samples (Figures S4,S5), though ENIGMA-OCD children showed moderate evidence linking greater symptom severity to stronger activation in the ACC (P+=0.94) and bilateral aI/fO (0.091<P+<0.93) (Figure 3B).

### Sensitivity analyses

Analyses restricted to stop-signal tasks in adults were largely consistent with the main results, but revealed stronger activation in medicated participants during response inhibition and weaker activation in late-onset OCD during error processing (Figures S6-S11). Leave-one-sample-out analyses for response inhibition showed generally consistent case-control and clinical subgroup results (Figure S13). However, when a large sample using an eye-blink suppression task (n=144) was excluded, evidence for case-control effects weakened further. Results were consistent across alternative analytic approaches, including ComBat harmonization (Figure S14B), modelling task instead of sample as a covariate (Figure S14C), and using a frequentist framework (Tables S5,S6).

## Discussion

In this mega-analysis we investigated neural activation differences in response inhibition and error processing between individuals with OCD and HCs, based on individual-level whole-brain activation maps from multiple inhibitory control tasks. Across ENIGMA-OCD and ABCD samples, inhibitory control tasks elicited robust activation in nearly all predefined ROIs associated with the inhibition subconstruct of the RDoC’s *cognitive systems* domain. Contrary to expectations, we did not find consistent evidence of weaker response inhibition or stronger error-processing activation in individuals with OCD or likely OCD in predefined regions. Instead, case-control differences during response inhibition varied across groups, as ABCD children showed reduced activation in the anterior insula/frontal operculum and right supplementary motor area, while ENIGMA-OCD adults and children showed greater activation in somatomotor and visual regions, respectively. In adults, OCD was also associated with weaker activation during error processing in multiple fronto-striatal regions, while ENIGMA-OCD children showed increased activation of the thalamus and frontoparietal and attention networks. Symptom severity was widely associated with brain activation, but in opposite directions in adult and pediatric OCD: greater symptom severity was correlated with weaker activation in adults but stronger activation in children. In contrast to previous ENIGMA-OCD analyses on structural and functional brain measures(26–28), there was little evidence for effects of medication status or age of onset on activation during inhibitory control.

Our whole-brain results align with Norman et al.’s meta-analysis combining adult and pediatric samples(4), as we observed increased activation in somatomotor regions in adults with OCD and in visual regions in children with OCD (ENIGMA-OCD samples) during response inhibition. This suggests prior studies may reflect the combined contributions of distinct effects in adult and pediatric samples, which we were able to disentangle by analyzing these samples separately. In contrast, our ROI analyses in the ENGIMA-OCD samples did not replicate earlier meta-analytic findings in adult and pediatric cohorts, such as weaker OCD activation in the frontal gyrus and anterior cingulate cortex(29), or the pattern reported by Norman et al. who observed weaker activation in frontoparietal and striatal regions but stronger activation in premotor, parietal, and temporal regions. This discrepancy may be due to ROIs being defined from peak activation in HCs, which may capture core regions of activation but miss peripheral differences that emerge in OCD when task demands increase.

During error processing, ENIGMA-OCD children with OCD showed increased error-related activation in the thalamus and frontoparietal and attention networks, consistent with earlier findings of increased fronto-insular and cingulate activity in a pediatric sample with sub-clinical OCD symptoms(30). In contrast, adults with OCD showed unexpectedly weaker activation in multiple fronto-striatal and attention regions, at odds with prior meta-analytic evidence of increased prefrontal and cingulate activity(4). This pattern may partly reflect symptom severity, which related to weaker activation in adults but stronger in children during error processing. However, this pattern conflicts with prior research, which has linked increased symptom severity to stronger error-related activity in an adult sample(31), but to weaker activity a pediatric sub-clinical sample(30).

Taken together, our findings support the idea that OCD follows distinct neurodevelopmental trajectories. Children and adults displayed opposite patterns of activation across the majority of brain regions during response inhibition - weaker in adults but stronger in children – with symptom severity further amplifying these divergent effects. This pattern is consistent with prior findings that children with OCD show stronger activation in prefrontal and anterior cingulate regions during response inhibition(32). Greater plasticity may allow children with OCD to compensate for diminished inhibitory control through stronger cortical activation to maintain performance. This pattern of heightened cortical recruitment is also seen in unaffected first-degree relatives of individuals with OCD during response inhibition(33) and error processing(34), and it may constitute a protective factor or trait-like neural marker in OCD. In contrast, adults’ hypoactivation suggests that by adulthood, altered maturation has reduced compensatory recruitment in inhibitory control networks and made them more sensitive to symptom states.

Consistent with our results, previous meta-analysis also did not find effects of illness duration or medication status on inhibitory control(4, 29). Although our large subject-level dataset improved statistical power to investigate clinical covariates, limited detail beyond medication status and age of onset may have constrained our ability to detect subgroup differences. Notably, medication effects emerged only in the ABCD sample, where medicated children with likely OCD showed stronger frontoparietal activation than their unmedicated peers. Because ABCD children were identified by parent-rated OCD symptoms rather than clinical evaluation, these findings require caution. Nonetheless, a concordance study reported 82% agreement between parent and clinician K-SADS reports for OCD diagnosis(35). Longitudinal studies are extremely sparse but are much needed to clarify how inhibitory control develops in OCD, and future follow-ups of the ABCD sample – even if some cases remain sub-clinical – offer a valuable opportunity to address this gap.

Our study relied on retrospective analysis of existing datasets collected on different scanners and task paradigms, capturing somewhat different cognitive processes. Findings were generally robust to sensitivity analyses, with the only notable shift occurring when excluding the sole eye-blink suppression task, suggesting that inhibition of involuntary autonomic responses engages different neural processes than voluntary motor behavior. The predominance of stop-signal tasks in our dataset may have biased results towards pure response inhibition (no alternative response) over cognitive inhibition (requiring substitution of an alternative response). This imbalance may partly explain inconsistent case-control effects, as a prior meta-analysis reported weaker activation in OCD specifically during cognitive inhibition tasks, but no overall differences when both domains were combined(36). Task demands are typically higher in response inhibition than in cognitive inhibition, and individuals with OCD often show hyperactivation when task demands are low, but hypoactivation when demands are high(37). Consistent with this, restricting our analyses to stop-signal tasks alone shifted the activation pattern from slightly increased to slightly decreased in OCD, though evidence for this was weak (Figure S8).

Nonetheless, the standardized processing protocol we applied across all datasets, the strong task effect we observed, and the robustness of our findings to sensitivity analyses suggest we successfully captured domain-level inhibitory control independent of individual task effects. While several meta-analyses have examined inhibitory control in OCD(4, 8–11, 29), our findings go beyond the previous literature by disentangling inhibitory control from broader executive function tasks, which previous studies conflated. Further, our results add to emerging evidence for distinct patterns of lifespan neurodevelopment and aging on neural recruitment during cognitive inhibition and response inhibition(38), underscoring the need to consider developmental stage and task demands when interpreting inhibitory control deficits in OCD.

## Conclusion

Our findings point to distinct effects of developmental stage and disease severity on neural activation during response inhibition and error processing. In childhood, inhibitory control activity in OCD is stronger, possibly reflecting compensatory processing, while in adulthood this reverses with weaker activation across key inhibitory network regions. These alterations were moderated by symptom severity, though other clinical factors had minimal influence on inhibitory activation, suggesting they may represent a symptom state.

## Acknowledgments

The ENIGMA-Obsessive Compulsive Disorder Working-Group gratefully acknowledges support from the International Obsessive-Compulsive Disorder Foundation with the Innovator Award of 2021 awarded to OAvdH and CV. The authors were further supported by the National Institute of Health (Grant No. R01MH138569 to PMT, OvdH, HBS; Grant Nos. R33MH107589 and R01MH111794 to ERS); the Dutch Research Council (Grant No. Vidi 917.15.318 to GvW); the European Union Horizon Seventh Framework Programme (Grant No. 278948 to JB); the Spanish Ministry of Science, Innovation and Universities (Grant No. ISCIII PI22/00752 to PA); the Catalan Audiovisual Media Corporation La Fundació Marató de TV3 (Grant No. 202201 to CSM and PA); the Carlos III Health Institute (Grant No. PI19/01179 to CSM); the German Research Foundation (Grant No. KO 3744/11-1 to KK and DRM); the German Center for Mental Health; the Western Norway Health Authority (Grants Nos. 911754 and 911880 to ALT); the Portuguese Foundation for Science and Technology (Grant No. UIDP/4950/2020 to MCB and ICD; Grant No. 2022.04701.PTDC to ICD); the Swiss National Science Foundation (Grant No. 320030_130237 to SW and SB); the Hartmann Müller Foundation (Grant No. 1460 to SB); the Carl Zeiss Foundation; the Alexander von Humboldt Foundation’s Alexander-von-Humboldt-Professorship award to Peter Dayan; the South African Medical Research Council; the Wellcome-Department of Biotechnology India Alliance (Grant No. 500236/Z/11/Z to GV); the Basic Research Program of the Korea Brain Research Institute and the National Research Foundation of Korea of the Korean Ministry of Science & Information and Communication Technology through the Brain Science Convergence Research Program (Grant Nos. RS-2023-00266120 and 25-BR-05-05 to MK).

Data used in the preparation of this article were obtained from the Adolescent Brain Cognitive Development^SM^ (ABCD) Study (https://abcdstudy.org), held in the NIMH Data Archive (NDA). This is a multisite, longitudinal study designed to recruit more than 10,000 children age 9-10 and follow them over 10 years into early adulthood. The ABCD Study® is supported by the National Institutes of Health and additional federal partners under award numbers U01DA041048, U01DA050989, U01DA051016, U01DA041022, U01DA051018, U01DA051037, U01DA050987, U01DA041174, U01DA041106, U01DA041117, U01DA041028, U01DA041134, U01DA050988, U01DA051039, U01DA041156, U01DA041025, U01DA041120, U01DA051038, U01DA041148, U01DA041093, U01DA041089, U24DA041123, U24DA041147. A full list of supporters is available at https://abcdstudy.org/federal-partners.html. A listing of participating sites and a complete listing of the study investigators can be found at https://abcdstudy.org/consortium_members/. ABCD consortium investigators designed and implemented the study and/or provided data but did not necessarily participate in the analysis or writing of this report. This manuscript reflects the views of the authors and may not reflect the opinions or views of the NIH or ABCD consortium investigators.

## Data availability

Processed and de-identified tabulated ROI-based and whole-brain parcellated brain activation data will be made available through the ENIGMA Toolbox (enigma-toolbox.readthedocs.io) in accordance with ENIGMA consortium policies. The code used for analyses is available at github.com/nadza-dz/task-based-fMRI-processing-pipeline-ENIGMA-OCD.git.

## References

1. Chambers CD, Garavan H, Bellgrove MA. Insights into the neural basis of response inhibition from cognitive and clinical neuroscience. Neurosci Biobehav Rev. 2009;33(5):631–46.

2. Insel T, Cuthbert B, Garvey M, Heinssen R, Pine SD, Quinn K, et al. Research Domain Criteria (RDoC): Toward a New Classification Framework for Research on Mental Disorders. American Journal of Psychiatry. 2010;167(7):748–51.

3. Shephard E, Stern ER, van den Heuvel OA, Costa DLC, Batistuzzo MC, Godoy PBG, et al. Toward a neurocircuit-based taxonomy to guide treatment of obsessive-compulsive disorder. Mol Psychiatry. 2021;26(9):4583–604.

4. Norman LJ, Taylor SF, Liu Y, Radua J, Chye Y, De Wit SJ, et al. Error Processing and Inhibitory Control in Obsessive-Compulsive Disorder: A Meta-analysis Using Statistical Parametric Maps. Biol Psychiatry. 2019;85(9):713–25.

5. van der Straten A, Bruin W, van de Mortel L, ten Doesschate F, Merkx MJM, de Koning P, et al. Pharmacological and Psychological Treatment Have Common and Specific Effects on Brain Activity in Obsessive-Compulsive Disorder. Depression and Anxiety. 2024;2024(1):6687657.

6. de Wit SJ, de Vries FE, van der Werf YD, Cath DC, Heslenfeld DJ, Veltman EM, et al. Presupplementary motor area hyperactivity during response inhibition: a candidate endophenotype of obsessive-compulsive disorder. Am J Psychiatry. 2012;169(10):1100–8.

7. Zhang J, Yang X, Yang Q. Neuropsychological dysfunction in adults with early-onset obsessive-compulsive disorder: the search for a cognitive endophenotype. Revista Brasileira de Psiquiatria. 2015;37(2):126–32.

8. Del Casale A, Rapinesi C, Kotzalidis GD, De Rossi P, Curto M, Janiri D, et al. Executive functions in obsessive-compulsive disorder: An activation likelihood estimate meta-analysis of fMRI studies. World J Biol Psychiatry. 2016;17(5):378–93.

9. Rasgon A, Lee WH, Leibu E, Laird A, Glahn D, Goodman W, et al. Neural correlates of affective and non-affective cognition in obsessive compulsive disorder: A meta-analysis of functional imaging studies. Eur Psychiatry. 2017;46:25–32.

10. Eng GK, Sim K, Chen S-HA. Meta-analytic investigations of structural grey matter, executive domain-related functional activations, and white matter diffusivity in obsessive compulsive disorder: An integrative review. Neuroscience & Biobehavioral Reviews. 2015;52:233–57.

11. Picó-Pérez M, Moreira PS, de Melo Ferreira V, Radua J, Mataix-Cols D, Sousa N, et al. Modality-specific overlaps in brain structure and function in obsessive-compulsive disorder: Multimodal meta-analysis of case-control MRI studies. Neuroscience & Biobehavioral Reviews. 2020;112:83–94.

12. van den Heuvel OA, Boedhoe PSW, Bertolin S, Bruin WB, Francks C, Ivanov I, et al. An overview of the first 5 years of the ENIGMA obsessive-compulsive disorder working group: The power of worldwide collaboration. Hum Brain Mapp. 2022;43(1):23–36.

13. Boedhoe PSW, Heymans MW, Schmaal L, Abe Y, Alonso P, Ameis SH, et al. An Empirical Comparison of Meta- and Mega-Analysis With Data From the ENIGMA Obsessive-Compulsive Disorder Working Group. Front Neuroinform. 2018;12:102.

14. Dzinalija N, Vriend C, Waller L, Simpson HB, Ivanov I, Agarwal SM, et al. Negative valence in Obsessive-Compulsive Disorder: A worldwide mega-analysis of task-based functional neuroimaging data of the ENIGMA-OCD consortium. Biological Psychiatry. 2024.

15. Dzinalija N, Veer IM, Simpson HB, Ivanov I, Balachander S, Benedetti F, et al. Executive control in Obsessive-Compulsive Disorder: A worldwide mega-analysis of task-based functional neuroimaging data of the ENIGMA-OCD consortium. bioRxiv. 2025:2025.08.20.671231.

16. Goodman KW. The Yale-Brown Obsessive Compulsive Scale. Archives of General Psychiatry. 1989;46(11):1006.

17. Scahill L, Riddle AM, Mcswiggin-Hardin M, Ort IS, King AR, Goodman KW, et al. Children’s Yale-Brown Obsessive Compulsive Scale: Reliability and Validity. Journal of the American Academy of Child & Adolescent Psychiatry. 1997;36(6):844–52.

18. Kaufman J, Birmaher B, Brent D, Rao UMA, Flynn C, Moreci P, et al. Schedule for Affective Disorders and Schizophrenia for School-Age Children-Present and Lifetime Version (K-SADS-PL): Initial Reliability and Validity Data. Journal of the American Academy of Child & Adolescent Psychiatry. 1997;36(7):980–8.

19. Kaufman J, Townsend LD, Kobak K. The Computerized Kiddie Schedule for Affective Disorders and Schizophrenia (KSADS): Development and Administration Guidelines. Journal of the American Academy of Child & Adolescent Psychiatry. 2017;56(10, Supplement):S357.

20. Waller L, Erk S, Pozzi E, Toenders YJ, Haswell CC, Büttner M, et al. ENIGMA HALFpipe: Interactive, reproducible, and efficient analysis for resting-state and task-based fMRI data. Hum Brain Mapp. 2022;43(9):2727–42.

21. Esteban O, Markiewicz JC, Blair WR, Moodie AC, Isik IA, Erramuzpe A, et al. fMRIPrep: a robust preprocessing pipeline for functional MRI. Nature Methods. 2019;16(1):111–6.

22. Chen G, Xiao Y, Taylor PA, Rajendra JK, Riggins T, Geng F, et al. Handling Multiplicity in Neuroimaging Through Bayesian Lenses with Multilevel Modeling. Neuroinformatics. 2019;17(4):515–45.

23. Johnson WE, Li C, Rabinovic A. Adjusting batch effects in microarray expression data using empirical Bayes methods. Biostatistics. 2007;8(1):118–27.

24. Schaefer A, Kong R, Gordon ME, Laumann OT, Zuo X-N, Holmes JA, et al. Local-Global Parcellation of the Human Cerebral Cortex from Intrinsic Functional Connectivity MRI. Cerebral Cortex. 2018;28(9):3095–114.

25. Tian Y, Margulies SD, Breakspear M, Zalesky A. Topographic organization of the human subcortex unveiled with functional connectivity gradients. Nature Neuroscience. 2020;23(11):1421–32.

26. Bruin WB, Taylor L, Thomas RM, Shock JP, Zhutovsky P, Abe Y, et al. Structural neuroimaging biomarkers for obsessive-compulsive disorder in the ENIGMA-OCD consortium: medication matters. Transl Psychiatry. 2020;10(1):342.

27. Ivanov I, Boedhoe PSW, Abe Y, Alonso P, Ameis SH, Arnold PD, et al. Associations of medication with subcortical morphology across the lifespan in OCD: Results from the international ENIGMA Consortium. Journal of Affective Disorders. 2022;318:204–16.

28. Bruin WB, Abe Y, Alonso P, Anticevic A, Backhausen LL, Balachander S, et al. The functional connectome in obsessive-compulsive disorder: resting-state mega-analysis and machine learning classification for the ENIGMA-OCD consortium. Molecular Psychiatry. 2023;28(10):4307–19.

29. Li Z, Tong G, Wang Y, Ruan H, Zheng Z, Cheng J, et al. Task fMRI studies investigating inhibitory control in patients with obsessive-compulsive disorder and eating disorders: A comparative meta-analysis. The World Journal of Biological Psychiatry. 2024;25(1):26–42.

30. Becker H, Liu Y, Hanna GL, Bilek E, Block SR, Hardee JE, et al. Error-related brain activity associated with obsessive–compulsive symptoms in youth. Brain and Behavior. 2023;13(4):e2941.

31. Agam Y, Greenberg JL, Isom M, Falkenstein MJ, Jenike E, Wilhelm S, et al. Aberrant error processing in relation to symptom severity in obsessive–compulsive disorder: A multimodal neuroimaging study. NeuroImage: Clinical. 2014;5:141–51.

32. Larsen KM, Uhre VF, Labanauskas V, Madsen KH, Nejad AB, Baaré W, et al. Inhibitory control in children and adolescents with paediatric-onset obsessive-compulsive disorder - An fMRI study. medRxiv. 2025:2025.05.09.25327300.

33. de Vries FE, de Wit SJ, Cath DC, van der Werf YD, van der Borden V, van Rossum TB, et al. Compensatory Frontoparietal Activity During Working Memory: An Endophenotype of Obsessive-Compulsive Disorder. Biological Psychiatry. 2014;76(11):878–87.

34. Grützmann R, Kaufmann C, Wudarczyk OA, Balzus L, Klawohn J, Riesel A, et al. Error-Related Brain Activity in Patients With Obsessive-Compulsive Disorder and Unaffected First-Degree Relatives: Evidence for Protective Patterns. Biological Psychiatry Global Open Science. 2022;2(1):79–87.

35. Townsend L, Kobak K, Kearney C, Milham M, Andreotti C, Escalera J, et al. Development of Three Web-Based Computerized Versions of the Kiddie Schedule for Affective Disorders and Schizophrenia Child Psychiatric Diagnostic Interview: Preliminary Validity Data. Journal of the American Academy of Child & Adolescent Psychiatry. 2020;59(2):309–25.

36. Funch Uhre V, Melissa Larsen K, Marc Herz D, Baaré W, Katrine Pagsberg A, Roman Siebner H. Inhibitory control in obsessive compulsive disorder: A systematic review and activation likelihood estimation meta-analysis of functional magnetic resonance imaging studies. NeuroImage: Clinical. 2022;36:103268.

37. van Velzen LS, de Wit SJ, Ćurĉić-Blake B, Cath DC, de Vries FE, Veltman DJ, et al. Altered inhibition-related frontolimbic connectivity in obsessive–compulsive disorder. Human Brain Mapping. 2015;36(10):4064–75.

38. Long J, Song X, Wang Y, Wang C, Huang R, Zhang R. Distinct neural activation patterns of age in subcomponents of inhibitory control: A fMRI meta-analysis. Front Aging Neurosci. 2022;14:938789.

