## Supplement for "Inhibitory control and error processing in Obsessive-Compulsive Disorder: A mega-analysis of task-based fMRI data by the ENIGMA-OCD consortium"

### Supplemental Methods

MRI acquisition parameters ........................................................ 2
Regions-of-interest coordinates ................................................ 4
Preprocessing and quality control guidelines............................ 5
Data exclusions ................................................................... 5
Missing brain activation data .................................................. 8
Explanation of RBA ridge plots ................................................. 11

### Supplemental Results

Demographics per site ............................................................ 12
Clinical covariate effects of response inhibition in adults ........ 13
Clinical covariate effects of error processing in adults ............ 15
Clinical covariate effects of response inhibition in children .... 16
Clinical covariate effects of error processing in children ......... 17
Clinical covariate effects of error processing in ABCD ............ 18
Main effect of response inhibition in SST tasks ........................ 19
Main effect of error processing in SST tasks ............................. 20
Case-control effect of response inhibition in SST tasks ........... 21
Case-control effect of error processing in SST tasks ................ 22
Clinical covariate effects of response inhibition in SST tasks ... 23
Clinical covariate effects of error processing in SST tasks ....... 24
Whole-brain Bayesian multilevel results .................................. 25
Sensitivity leave-one-sample-out analyses ............................... 26
Sensitivity sample and task correction methods ...................... 31
Sensitivity frequentist statistics ................................................ 32

Supplemental references ........................................................ 35

### Supplemental Methods

**Table S1 –** MRI acquisition parameters of studies included in mega-analysis

| Site-PI | Scanner | Head coil (# channels) | Pulse sequence | Single-/ multiband | Matrix | Volumes (N) | Slices (N) | Scan order | Slice gap (mm) | TR (ms) | TE (ms) | Flip angle (°) | FOV (mm) | Voxel size (mm) | Slice time correction applied | Prospective motion correction during scanning |
|---|---|---|---|---|---|---|---|---|---|---|---|---|---|---|---|---|
| **van den Heuvel (1)** | 3T GE Signa HDxt | 8 | GE-EPI | singleband | 64x64 | ~430 | 40 | ascending | 0.2 | 2100 | 30 | 80 | 240 | 3mm$^3$ | Yes | No |
| **van den Heuvel (2)** | 3T GE Discovery MR750 | 32 | GE-EPI | singleband | 64x64 | 307 | 42 | ascending | 0.33 | 2200 | 28 | 90 | 211 | 3.3x3.3x3 | Yes | No |
| **van Wingen (3)** | 3T Philips Achieva | 32 | GE-EPI | singleband | 75x75 | 197 | 37 | | 0.3 | 2375 | 9,26,44 | 76 | 224x224 | 2.8x2.8x3 | No | No |
| **Huyser (4)** | 3T Philip Intera MR | 6 | GE-EPI | singleband | 96x96 | 250 | 40 | interleaved | 0 | 2300 | 30 | 90 | 220x120x220 | 2.29x2.29x3 | No | No |
| **Reddy** | 3T Siemens Skyra | 32 | SS-EPI | singleband | 64x64 | 126 | 37 | descending | 0.8 | 2000 | 30 | 78 | 192 | 3mm$^3$ | Yes | No |
| **Menchón / Soriano-Mas (5)** | 1.5T GE Signa Excite | 8 | GE-EPI | singleband | 64x64 | 180 | 22 | interleaved | 1 | 2000 | 50 | 90 | 240 | 3.75x3.75x4 | Yes | No |
| **Menchón / Soriano-Mas** | 3T Philips Ingenia | 32 | SS-EPI | singleband | 80x80 | 97 | 40 | interleaved | 0 | 2000 | 25 | 90 | 240 | 3mm$^3$ | Yes | No |
| **Fullana** | 3T Philips Ingenia | 32 | GE-EPI | singleband | 96x96 | 100 | 32 | interleaved | 0.75 | 2000 | 30 | 70 | 240x240 | 2.5x2.5x4.25 | Yes | No |
| **Thorsen** | 3T GE Discovery MR750 | 8 | GE-EPI | singleband | 64x64 | 430 | 34 | interleaved | 0.2 | 2100 | 30 | 80 | 220 | 3.44x3.44x3 | Yes | No |
| **Beucke / Kathmann (6)** | 3T Siemens Magneton Trio Tim MR | 32 | GE-EPI | singleband | 64x64 | 720 | 32 | descending | 25% | 1940 | 30 | 78 | 192 | 3x3x3.75 | Yes | No |

| | | | | | | | | | | | | | | | | |
|---|---|---|---|---|---|---|---|---|---|---|---|---|---|---|---|---|
| **Castelo-Branco** | 3T Siemens Prisma | 64 | CMRR EPI | 6 | 100x100 | ~360 | 37 | interleaved | 0 | 1000 | 37 | 68 | 200x200 | 2mm$^3$ | Yes | No |
| **Koch** | 3T Philips Ingenia | 32 | GE-EPI | 2 | 64x64 | 660 | 36 | interleaved | 0.3 | 1000 | 30 | 60 | 100 | 3.0x3.0x3.75 | No | No |
| **Stern (7)** | 3T Siemens NKI TRIO | 32 | CMRR EPI 2D | 6 | 108x108 | 440 x 2 runs | 72 | interleaved | 0 | 1000 | 25.4 | 60 | 228 | 2.1mm$^3$ | No | No |
| **Buitelaar / van Rooij (8)** | 3T Siemens Prismafit | 12 | SMS-EPI | singleband | 64 | 263 | 36 | interleaved | 0 | 2100 | 35 | 74 | 192 | 3mm$^3$ | No | No |
| **Kwon (9)** | 1.5T Siemens Avanto | 12 | GE-EPI | singleband | 64x64 | 486 | 25 | interleaved | 0 | 2340 | 52 | 90 | 220 | 3.44x3.44x5 | No | No |
| **Walitza / Brem (10)** | 3T Philipps Achieva | 32 | GE-EPI | singleband | 96x96 | 333 x 2 runs | 40 | ascending | 0.7 | 1850 | 20 | 85 | 240 | 2.5mm$^3$ | Yes | No |
| | 3T Siemens Prisma | 32/64 | GE-EPI | 6 | 90x90 | ~437 x 2 runs | 60 | interleaved | 0 | 800 | 30 | 52 | 216x216 | 2.4mm$^3$ | No | No |
| | 3T Philips Achieva, Ingenia | 32 | GE-EPI | 6 | 90x90 | ~437 x 2 runs | 60 | interleaved | 0 | 800 | 30 | 52 | 216x216 | 2.4mm$^3$ | No | No |
| **ABCD sample *** | 3T GE Discovery MR750, DV25-26 | 32 | GE-EPI | 6 | 90x90 | ~437 x 2 runs | 60 | interleaved | 0 | 800 | 30 | 52 | 216x216 | 2.4mm$^3$ | No | No |

* ABCD samples were acquired at 18 different sites, on one of three MRI scanners (Siemens, Philips, or GE) with harmonized acquisition parameters

**Table S2 -** Regions of interest identified from Norman et al. (2019;(11)) showing whole-brain activation in healthy controls during inhibitory control tasks for the response inhibition and error processing contrasts. Coordinates are in MNI152 NLIN 6th generation space. ACC = anterior cingulate cortex; aI/fO = anterior insula/frontal operculum; dlPFC = dorsolateral prefrontal cortex; Occ = occipital; PMC = primary motor cortex; SMA = supplementary motor area; SMG = supramarginal gyrus; SPL = superior parietal lobule; STG = superior temporal gyrus.

| **Region** | **Lateralization** | **MNI coordinates** | | |
|---|---|---|---|---|
| | | **x** | **y** | **z** |
| **Response inhibition [stop success > go]** | | | | |
| aI/fO | right | 34 | 18 | 8 |
| | left | -40 | 10 | 2 |
| SMA | right | 4 | 14 | 58 |
| | left | -4 | 14 | 58 |
| SPL | right | 42 | -44 | 58 |
| | left | -42 | -44 | 58 |
| Occ. lobe | right | 28 | -72 | 36 |
| | left | -28 | -72 | 36 |
| PMC | right | 38 | -8 | 62 |
| | left | -48 | 2 | 24 |
| dlPFC | right | 34 | 44 | 34 |
| | left | -34 | 44 | 34 |
| **Error processing [stop fail > stop success]** | | | | |
| aI/fO | right | 44 | 14 | 2 |
| | left | -48 | 16 | -2 |
| STG | left | -40 | -8 | -12 |
| ACC | bilateral | 0 | 34 | 32 |
| Occ. lobe | bilateral | 0 | -82 | 18 |
| SMG | right | 66 | -40 | 28 |
| | left | -58 | -46 | 30 |

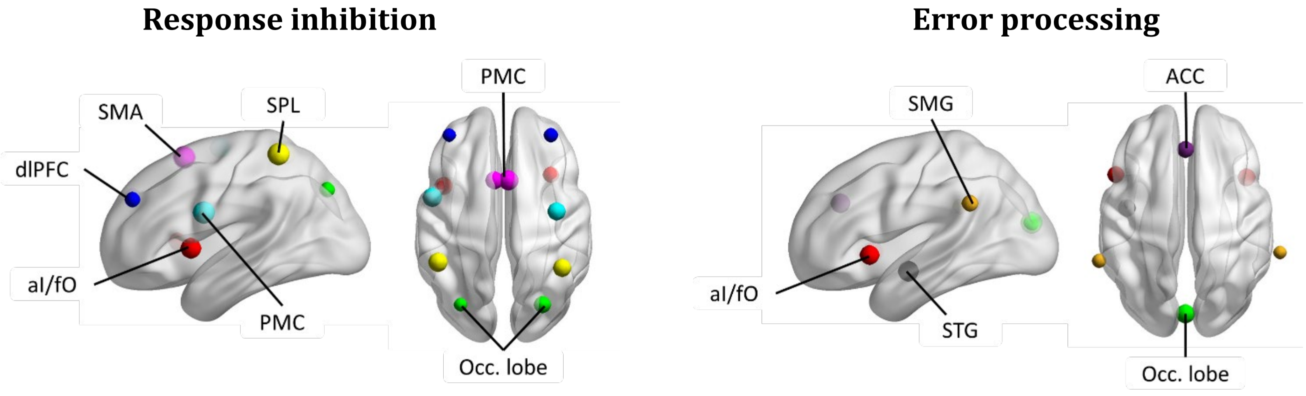

#### Preprocessing and quality control guidelines

Preprocessing was done at each site according to harmonized guidelines and each site performed quality-control on their own preprocessed data following harmonized guidelines which can be retrieved from the online repository doi.org/10.5281/zenodo.17141947.

#### Data exclusions

For twenty (20) participants (2 Obsessive-Compulsive Disorder [OCD] participants and 18 healthy control [HC] participants) the motion-correction denoising portion of the analysis pipeline failed to run in the error processing contrast only, sixty-four (64) participants (39 OCD and 25 HCs) failed quality assessment due to poor skull-stripping, registration or normalization, and sixty-one (61) participants (44 OCD and 17 HCs) were excluded due to excessive motion.

*Adult samples*

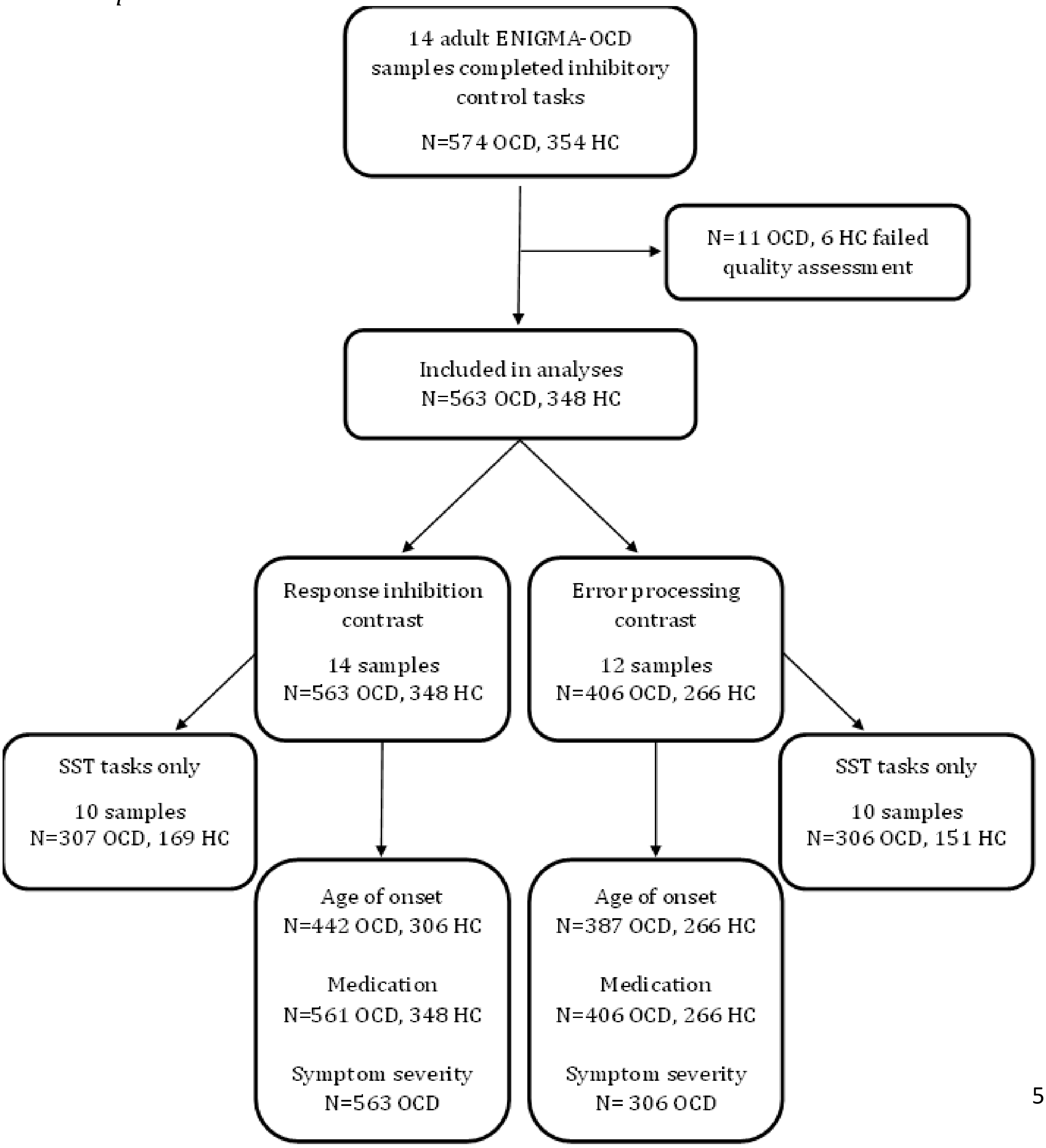

*Pediatric samples*

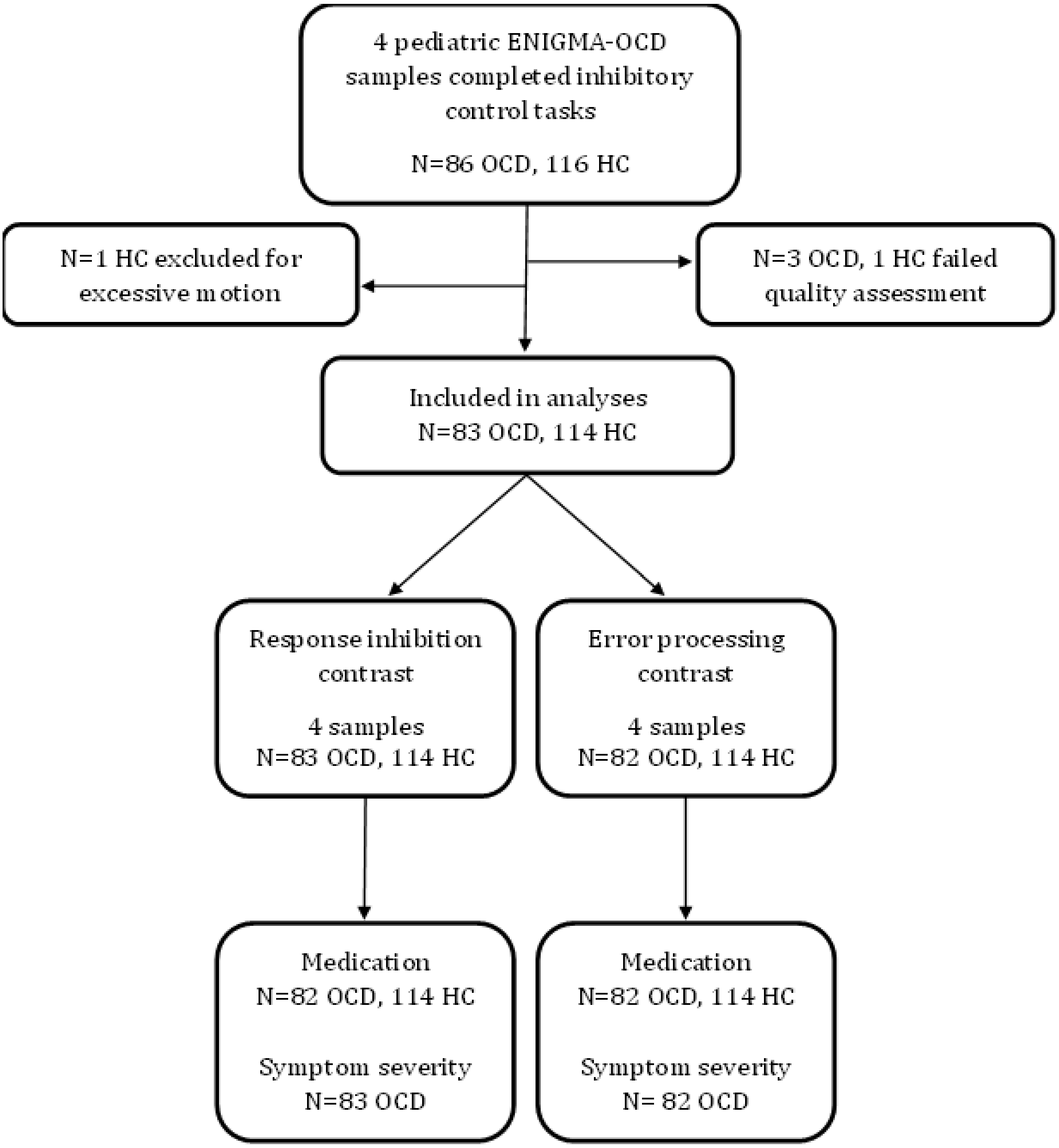

*ABCD sample*

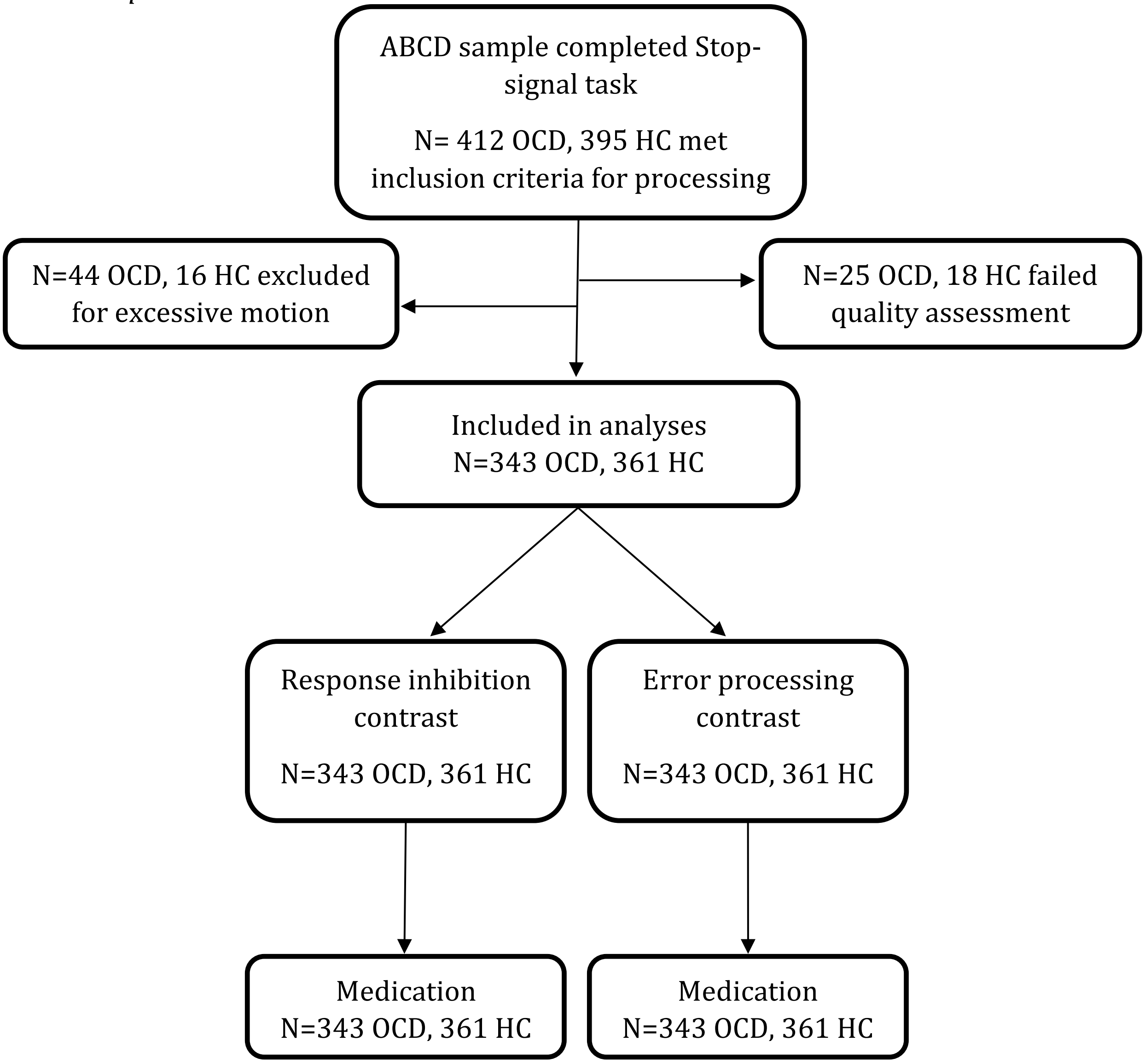

Clinical and demographic data from the Adolescent Brain Cognitive Development (ABCD) study were obtained from the 4.0 Curated Data Release (DOI: 10.15154/1523041). Raw imaging data were accessed from the ABCD Fast Track data release and are available via the NIMH Data Archive (https://nda.nih.gov/edit_collection.html?id=2573). We selected participants with likely OCD based on parent-reported diagnoses in the computerized KSADS interview. For each OCD participant, one healthy control (HC) was matched on data collection site, age, sex, and socioeconomic status. HCs were required to have no diagnosed or parent-reported psychiatric or developmental disorders and no psychoactive medication use. To ensure data quality and task engagement, participants were required to meet minimum performance criteria on the stop-signal task: at least 60% correct Go trials, at least 25% and no more than 75% correct Stop trials. Only participants with complete imaging and behavioral data were included, defined as having a T1-weighted scan, BOLD data from at least one run, field maps, behavioral log files, and relevant demographic information.

### Missing brain activation data

Brain activation data were included in ROI or whole-brain parcel analyses only if at least 30% of voxels in the ROI or parcel contained signal. ROIs or parcels not meeting this threshold were excluded; missing values were not imputed. No ROIs failed this criterion for either response inhibition or error processing contrasts in any cohort (ENIGMA-OCD adults, ENIGMA-OCD children, ABCD children). In whole-brain analyses, some parcels near the edges of the brain had insufficient coverage across contrasts and cohorts.

**Table S3 – Whole-brain parcels missing data per analysis, ordered by percent of sample missing data in given parcel** L = left hemisphere; OFC = orbitofrontal cortex; PFC = prefrontal cortex; PFCdPFCm = dorsomedial prefrontal cortex; PFCl = lateral prefrontal cortex; Post = posterior; R = right hemisphere; Temp = temporal; TempOccPar = temporo-occipito-parietal; VentroAnt = ventroanterior; VentroPost = ventroposterior.

| **ENIGMA-OCD adult samples** | | **ENIGMA-OCD pediatric samples** | | **ABCD sample** | |
|---|---|---|---|---|---|
| Parcel | % missing | Parcel | % missing | Parcel | % missing |
| **Response Inhibition** | | | | | |
| Limbic OFC 1 R | 24.6 | Limbic OFC 1 R | 7.1 | Limbic OFC 1 R | 6.0 |
| Limbic OFC 2 L | 21.9 | Limbic TempPole 1 L | 6.6 | Visual 5 L | 5.8 |
| Limbic TempPole 1 R | 13.6 | Visual 5 L | 6.1 | Limbic OFC 2 L | 2.3 |
| Limbic TempPole 1 L | 13.2 | Limbic TempPole 1 R | 5.6 | Limbic TempPole 2 R | 2.3 |
| Limbic TempPole 2 R | 11.4 | Limbic TempPole 2 R | 4.6 | Default Temp 2 L | 1.6 |
| Limbic TempPole 3 L | 10.1 | Limbic TempPole 3 L | 3.6 | Limbic TempPole 1 L | 0.7 |
| Default Temp 2 L | 7.2 | Limbic OFC 2 L | 3.0 | Limbic TempPole 2 L | 0.7 |
| Limbic TempPole 2 L | 5.7 | Default Temp 1 L | 2.0 | Visual 8 R | 0.7 |
| Limbic TempPole 3 R | 3.4 | Limbic OFC 3 R | 1.5 | Limbic OFC 3 R | 0.6 |
| Visual 5 L | 2.6 | Limbic TempPole 2 L | 1.5 | Control PFCl 1 R | 0.4 |
| Limbic OFC 3 R | 2.4 | Limbic TempPole 3 R | 1.5 | Limbic TempPole 3 L | 0.3 |
| Default Temp 1 L | 2.0 | Visual 8 R | 1.5 | Limbic TempPole 3 R | 0.3 |
| Control PFCl 1 R | 1.3 | Control PFCl 1 L | 1.0 | Control OFC 1 L | 0.1 |
| Default Temp 2 R | 1.2 | Control Temp 1 L | 1.0 | Control PFCl 1 L | 0.1 |
| Default PFC 4 L | 1.1 | Default Temp 2 L | 1.0 | Default Temp 1 L | 0.1 |
| Default Temp 1 R | 1.1 | Control OFC 1 L | 0.5 | Default Temp 1 R | 0.1 |
| Control Temp 1 L | 0.8 | Control PFCl 2 R | 0.5 | Limbic TempPole 1 R | 0.1 |
| Control PFCl 1 L | 0.7 | Default PFC 4 L | 0.5 | Visual 14 R | 0.1 |
| Default PFCdPFCm 1 R | 0.7 | Visual 11 L | 0.5 | Visual 6 R | 0.1 |
| Control OFC 1 L | 0.5 | Visual 2 L | 0.5 | Visual 9 L | 0.1 |
| Visual 6 R | 0.4 | Visual 3 L | 0.5 | | |
| Visual 8 R | 0.4 | Visual 6 R | 0.5 | | |
| Limbic OFC 1 L | 0.3 | Visual 7 L | 0.5 | | |
| Visual 7 L | 0.3 | Visual 9 L | 0.5 | | |
| Control PFCl 2 L | 0.2 | | | | |

| | | | | | |
|---|---|---|---|---|---|
| Control PFCl 2 R | 0.2 | | | | |
| Control Temp 1 R | 0.2 | | | | |
| Default PFC 2 L | 0.2 | | | | |
| Limbic OFC 2 R | 0.2 | | | | |
| Visual 9 L | 0.2 | | | | |
| Default PFC 13 L | 0.1 | | | | |
| Default PFC 3 L | 0.1 | | | | |
| Default Temp 3 L | 0.1 | | | | |
| Dorsal Attention Post 1 L | 0.1 | | | | |
| Dorsal Attention Post 10 L | 0.1 | | | | |
| Dorsal Attention Post 10 R | 0.1 | | | | |
| Visual 1 R | 0.1 | | | | |
| Visual 2 L | 0.1 | | | | |
| **Error Processing** | | | | | |
| Limbic OFC 1 R | 30.2 | Limbic OFC 1 R | 7.1 | Limbic OFC 1 R | 6.0 |
| Limbic OFC 2 L | 28.0 | Limbic TempPole 1 L | 6.6 | Visual 5 L | 5.8 |
| Limbic TempPole 1 R | 15.1 | Visual 5 L | 5.6 | Limbic OFC 2 L | 2.3 |
| Limbic TempPole 1 L | 14.1 | Limbic TempPole 1 R | 5.1 | Limbic TempPole 2 R | 2.3 |
| Limbic TempPole 2 R | 12.7 | Limbic TempPole 2 R | 4.6 | Default Temp 2 L | 1.6 |
| Limbic TempPole 3 L | 10.4 | Limbic TempPole 3 L | 3.6 | Limbic TempPole 1 L | 0.7 |
| Default Temp 2 L | 8.7 | Limbic OFC 2 L | 3.1 | Limbic TempPole 2 L | 0.7 |
| Limbic TempPole 2 L | 7.0 | Default Temp 1 L | 1.5 | Visual 8 R | 0.7 |
| Limbic TempPole 3 R | 3.9 | Limbic OFC 3 R | 1.5 | Limbic OFC 3 R | 0.6 |
| Visual 5 L | 3.0 | Limbic TempPole 2 L | 1.5 | Control PFCl 1 R | 0.4 |
| Limbic OFC 3 R | 2.7 | Limbic TempPole 3 R | 1.5 | Limbic TempPole 3 L | 0.3 |
| Default Temp 1 L | 2.1 | Visual 8 R | 1.5 | Limbic TempPole 3 R | 0.3 |
| Control PFCl 1 R | 1.8 | Control PFCl 1 L | 1.0 | Control OFC 1 L | 0.1 |
| Default Temp 1 R | 1.5 | Control Temp 1 L | 1.0 | Control PFCl 1 L | 0.1 |
| Default Temp 2 R | 1.5 | Default Temp 2 L | 1.0 | Default Temp 1 L | 0.1 |
| Default PFC 4 L | 1.3 | Control OFC 1 L | 0.5 | Default Temp 1 R | 0.1 |
| Control Temp 1 L | 0.9 | Control PFCl 2 R | 0.5 | Limbic TempPole 1 R | 0.1 |
| Default PFCdPFCm 1 R | 0.9 | Default PFC 4 L | 0.5 | Visual 14 R | 0.1 |
| Control OFC 1 L | 0.7 | Visual 11 L | 0.5 | Visual 6 R | 0.1 |
| Control PFCl 1 L | 0.7 | Visual 2 L | 0.5 | Visual 9 L | 0.1 |
| Limbic OFC 1 L | 0.4 | Visual 3 L | 0.5 | | |
| Visual 6 R | 0.4 | Visual 6 R | 0.5 | | |
| Visual 7 L | 0.4 | Visual 7 L | 0.5 | | |
| Control PFCl 2 L | 0.3 | Visual 9 L | 0.5 | | |
| Control PFCl 2 R | 0.3 | | | | |
| Control Temp 1 R | 0.3 | | | | |
| Default PFC 2 L | 0.3 | | | | |
| Limbic OFC 2 R | 0.3 | | | | |
| Visual 8 R | 0.3 | | | | |
| Default PFC 13 L | 0.1 | | | | |

| | |
|---|---|
| Default PFC 3 L | 0.1 |
| Default Temp 3 L | 0.1 |
| Dorsal Attention Post 1 L | 0.1 |
| Dorsal Attention Post 10 L | 0.1 |
| Dorsal Attention Post 10 R | 0.1 |
| Visual 1 R | 0.1 |
| Visual 2 L | 0.1 |
| Visual 9 L | 0.1 |

**Explanation of how to interpret ridge plot results of Regional Bayesian Analysis**

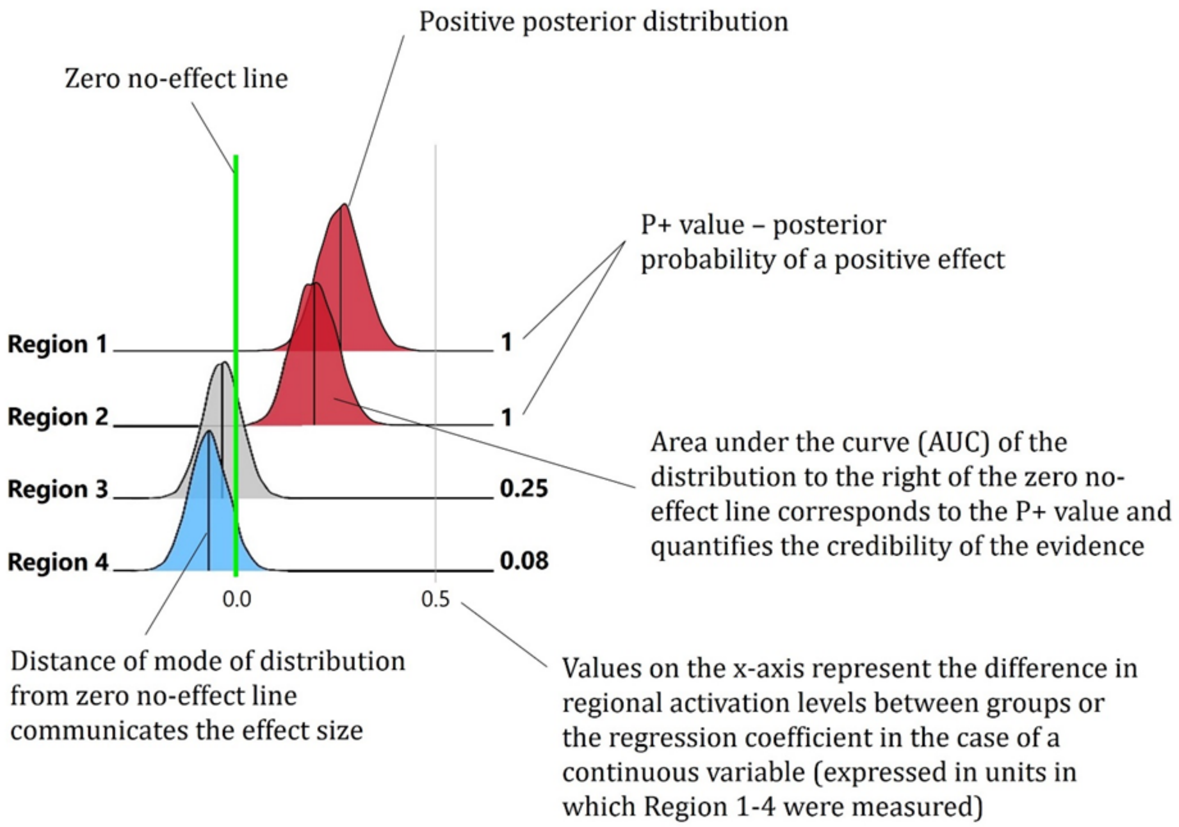

Consider this hypothetical analysis of effects in four regions of interest as performed under Regional Bayesian Analysis (RBA; Chen et al., 2019 (12)). In Regions 1 and 2 there is very strong evidence of a positive effect as the entire AUC is to the right of the no-effect line (P+=1.0), denoting a nearly 100% probability of the effect being positive in these regions. Region 1 has a larger effect size than Region 2 as the mode of the distribution is further away from the no-effect line. In Region 4 there is moderate evidence for a negative effect as AUC is largely to the left of the no-effect line (P+=0.08). Small values of P+ convey evidence that the effect is negative – a P+ value of 0.08 indicates that the probability of the effect being positive is only 8%, so the probability of it being negative is 92%. For Region 3, the distribution crosses the no-effect line with 25% of the AUC to the right of the no effect line (P+=0.25), meaning there is no strong evidence of a positive effect, but also no strong evidence of a negative effect in this region.

### Supplemental Results

#### Table S4 - Demographics per site

| Site-PI | City, Country, Sample | n OCD | n HC | % Male | Age (mean ± SD) | % Child onset OCD | % Medicated OCD | (C)Y-BOCS (mean ± SD) |
|---|---|---|---|---|---|---|---|---|
| **Adult ENIGMA-OCD samples** | | | | | | | | |
| van den Heuvel | Amsterdam, NLD I | 40 | 32 | 51.39 | 38.75 ± 10.29 | 62.50 | 0 | 21.8 ± 6.12 |
| van den Heuvel | Amsterdam, NLD II | 67 | 16 | 34.94 | 35.66 ± 12.11 | 56.72 | 62.69 | 28.41 ± 4.8 |
| van Wingen | Amsterdam, NLD III | 29 | 15 | 52.27 | 29.82 ± 8.65 | 58.62 | 55.17 | 25.86 ± 3.67 |
| Reddy | Bangalore, IND | 53 | 22 | 53.33 | 28.15 ± 5.75 | 30.19 | 35.85 | 24.51 ± 7.16 |
| Menchón / Soriano-Mas | Barcelona, ESP I | 22 | 0 | 45.45 | 35.59 ± 9.94 | 59.09 | 100 | 24.86 ± 6.47 |
| Menchón / Soriano-Mas | Barcelona, ESP II | 17 | 18 | 54.29 | 45.63 ± 8.83 | 47.06 | 94.12 | 23.41 ± 5.99 |
| Fullana | Barcelona, ESP III | 13 | 18 | 41.94 | 29.23 ± 8.49 | 76.92 | 0 | 24 ± 5.07 |
| Thorsen | Bergen, NOR | 38 | 30 | 38.24 | 30.19 ± 9.38 | 39.47 | 23.68 | 26.74 ± 4.38 |
| Beucke / Kathmann | Berlin, GER | 84 | 98 | 43.96 | 31.59 ± 9.55 | 60.71 | 41.67 | 22.17 ± 5.00 |
| Castelo-Branco | Coimbra, PRT | 19 | 22 | 100 | 28.46 ± 7.97 | 78.95 | 94.74 | 24.89 ± 5.84 |
| Koch | Munich, GER | 47 | 0 | 36.17 | 32.19 ± 11.91 | 68.09 | 65.96 | 19.9 ± 5.51 |
| Stern | New York, USA | 102 | 42 | 36.81 | 30.17 ± 11.42 | 0 | 54.90 | 23.3 ± 5.19 |
| Kwon | Seoul, KOR | 15 | 18 | 60.61 | 25.36 ± 4.03 | 40.00 | 0 | 21.07 ± 7.63 |
| Walitza / Brem | Zurich, CHE | 17 | 17 | 38.24 | 31.49 ± 8.36 | 35.29 | 58.82 | 17.12 ± 9.94 |
| **Pediatric ENIGMA-OCD samples** | | | | | | | | |
| Huyser | Amsterdam, NLD IV | 30 | 24 | 38.89 | 13.33 ± 2.58 | 100 | 0 | 24.86 ± 4.95 |
| Fullana | Barcelona, ESP III | 25 | 19 | 52.27 | 13.64 ± 2.43 | 100 | 0 | 25.09 ± 4.56 |
| Buitelaar / van Rooij | Nijmegen, NLD | 13 | 52 | 70.77 | 10.78 ± 1.08 | 100 | 23.08 | 18.23 ± 5.37 |
| Walitza / Brem | Zurich, CHE | 15 | 19 | 64.71 | 15.04 ± 1.48 | 100 | 53.33 | 16.92 ± 9.68 |
| **Pediatric ABCD sample** | | | | | | | | |
| ABCD study | USA | 343 | 361 | 53.41 | 9.98 ± 0.59 | 100 | 17.49 | - |

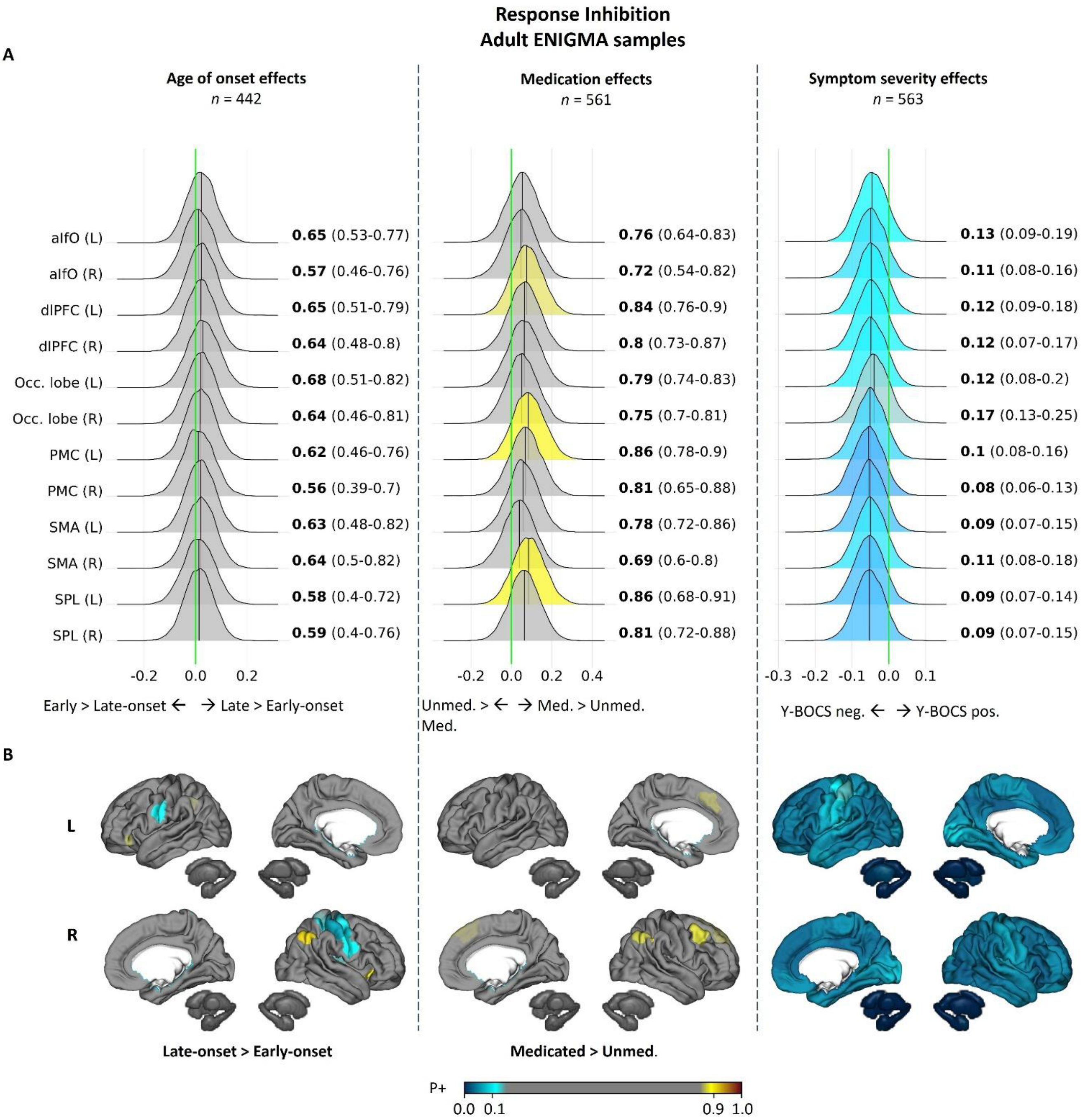

***Figure S1 - Effects of clinical features of OCD on response inhibition in adults from the ENIGMA-OCD sample*** (A) Region-of-interest results from Bayesian multilevel analyses of the effects of age of onset, medication status, and symptom severity during response inhibition. Posterior probability distributions express the credibility of an effect in each region. Next to each distribution the posterior probability of a positive effect (P+) is shown in bold, as well as the range of values this probability took on in leave-one-sample-out sensitivity analyses. In regions with posterior distributions to the right of the green no-effect line there is evidence of stronger activation during response inhibition while in regions with posterior distributions to the left of this line there is evidence of weaker activation during response inhibition. Regions are color-coded to reflect the strength of evidence for an effect, where in (darker) red regions there is stronger evidence of activation during response inhibition (P+ values >0.90 indicate moderate to very strong evidence for a positive effect). In (darker) blue regions there is stronger evidence of deactivation during response inhibition (P+ values <0.10 indicate moderate to very strong evidence for a negative effect). In grey regions there is no strong evidence of activation or deactivation during response inhibition. Values on the x-axis represent the difference in regional activation levels between the investigated groups (expressed as difference in Z-scores). (B) Whole-brain effects from Bayesian multilevel analyses. P+ values derived from Bayesian multilevel analyses denote the probability that there is stronger brain activation in a given region of the Schaefer 200-parcel 7-network cortical atlas and Melbourne 32-region subcortical atlas during response inhibition. Displayed are lateral and medial views of the cortex and subcortex. aI/fO = anterior insula/frontal operculum; dlPFC = dorsolateral prefrontal cortex; L = left; Occ = occipital; PMC = primary motor cortex; R = right; SMA = supplementary motor area; SPL = superior parietal lobule; Y-BOCS = Yale-Brown Obsessive-Compulsive Scale.

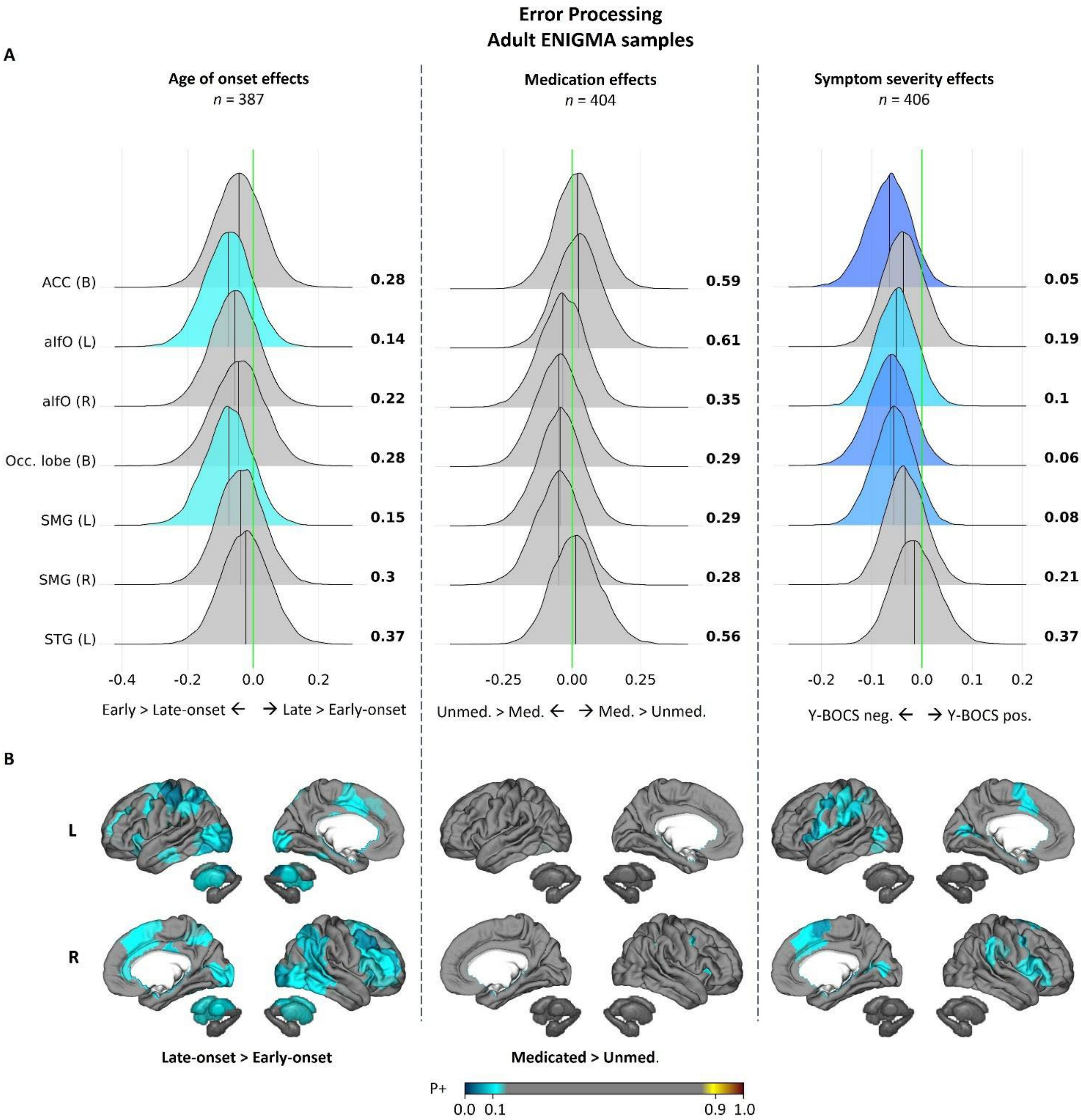

***Figure S2- Effects of clinical features of OCD on error processing in adults from the ENIGMA-OCD sample*** (A) Region-of-interest effects of age of onset, medication status, and symptom severity on activation during error processing. (B) Whole-brain analyses of error processing. ACC = anterior cingulate cortex; aI/fO = anterior insula/frontal operculum; L = left; Occ = occipital; R = right; SMG = supramarginal gyrus; STG = superior temporal gyrus; Y-BOCS = Yale-Brown Obsessive-Compulsive Scale.

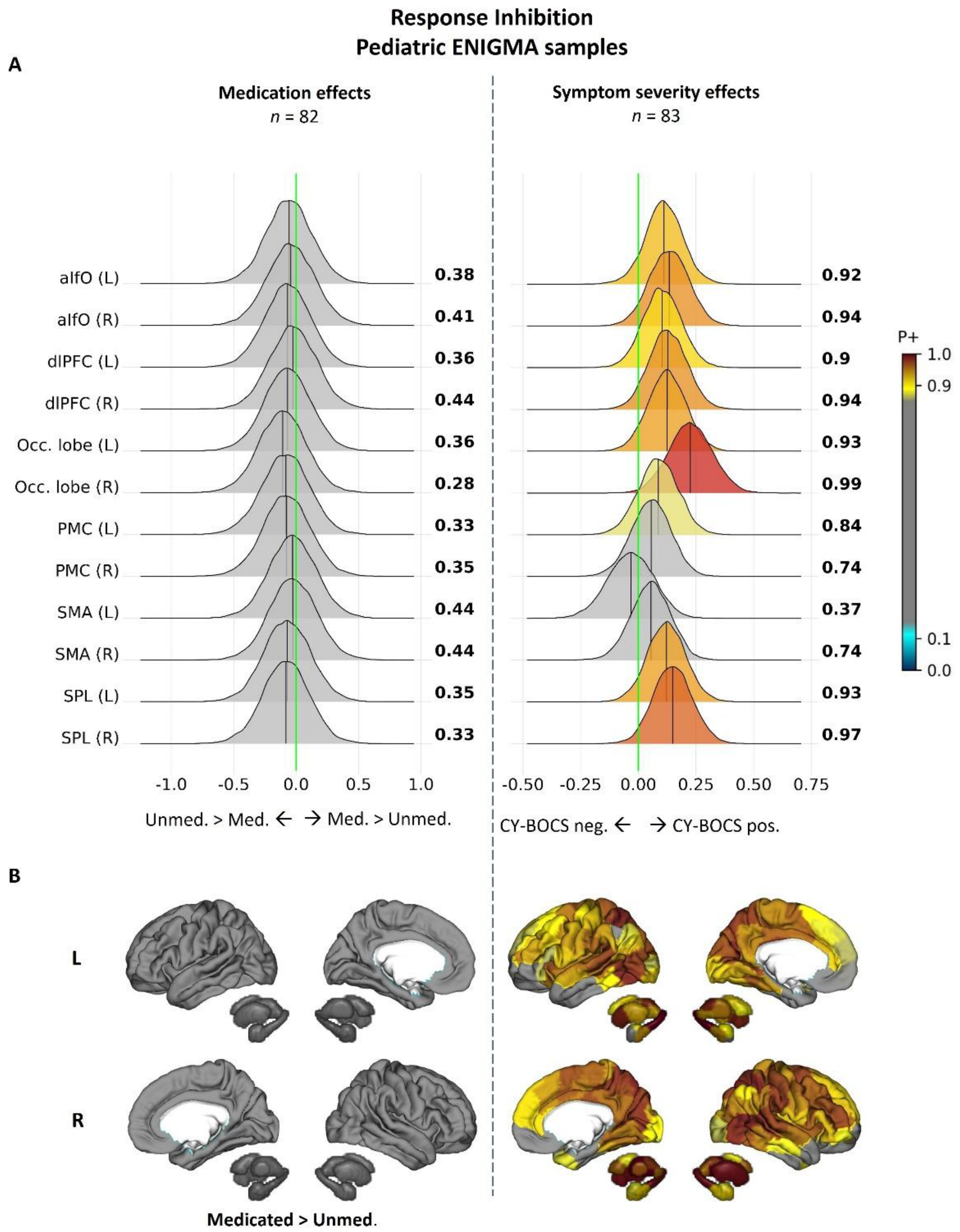

***Figure S3 - Effects of clinical features of OCD on response inhibition in children from the ENIGMA-OCD sample*** (A) Region-of-interest effects of medication status and symptom severity on activation during response inhibition. (B) Whole-brain analyses of response inhibition. aI/fO = anterior insula/frontal operculum; CY-BOCS = Children's Yale-Brown Obsessive-Compulsive Scale; dlPFC = dorsolateral prefrontal cortex; L = left; Occ = occipital; PMC = primary motor cortex; R = right; SMA = supplementary motor area; SPL = superior parietal lobule.

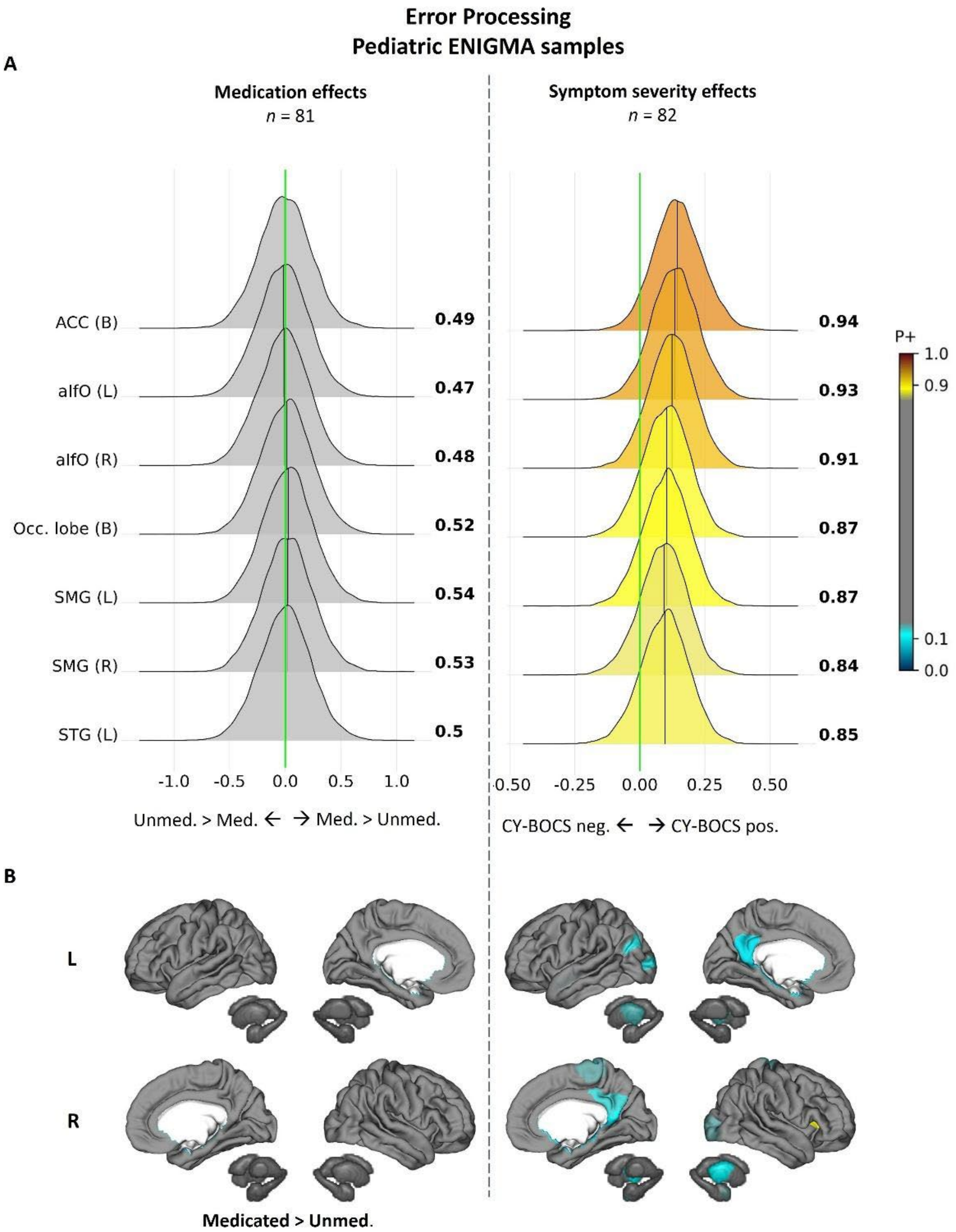

***Figure S4 - Effects of clinical features of OCD on error processing in children from the ENIGMA-OCD sample*** (A) Region-of-interest effects of medication status and symptom severity on activation during error processing. (B) Whole-brain analyses of error processing. ACC = anterior cingulate cortex; aI/fO = anterior insula/frontal operculum; CY-BOCS = Children's Yale-Brown Obsessive-Compulsive Scale; L = left; Occ = occipital; R = right; SMG = supramarginal gyrus; STG = superior temporal gyrus.

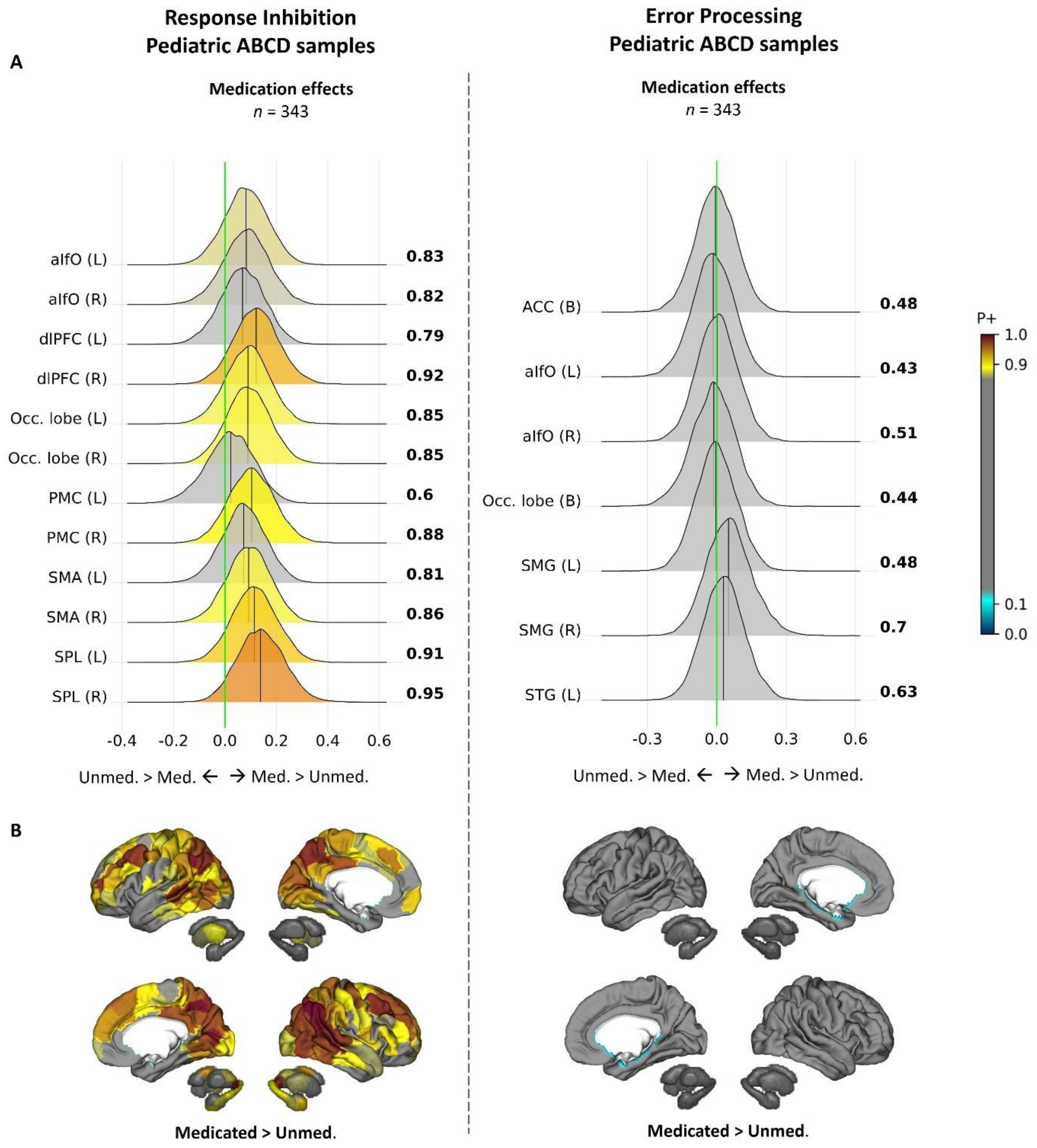

***Figure S5 - Effects of medication on response inhibition and error processing in children from the ABCD sample*** (A) Region-of-interest effects of medication status on activation during response inhibition and error processing. (B) Whole-brain analyses of response inhibition and error processing. ACC = anterior cingulate cortex; aI/fO = anterior insula/frontal operculum; dlPFC = dorsolateral prefrontal cortex; L = left; Occ = occipital; PMC = primary motor cortex; R = right; SMA = supplementary motor area; SMG = supramarginal gyrus; SPL = superior parietal lobule; STG = superior temporal gyrus.

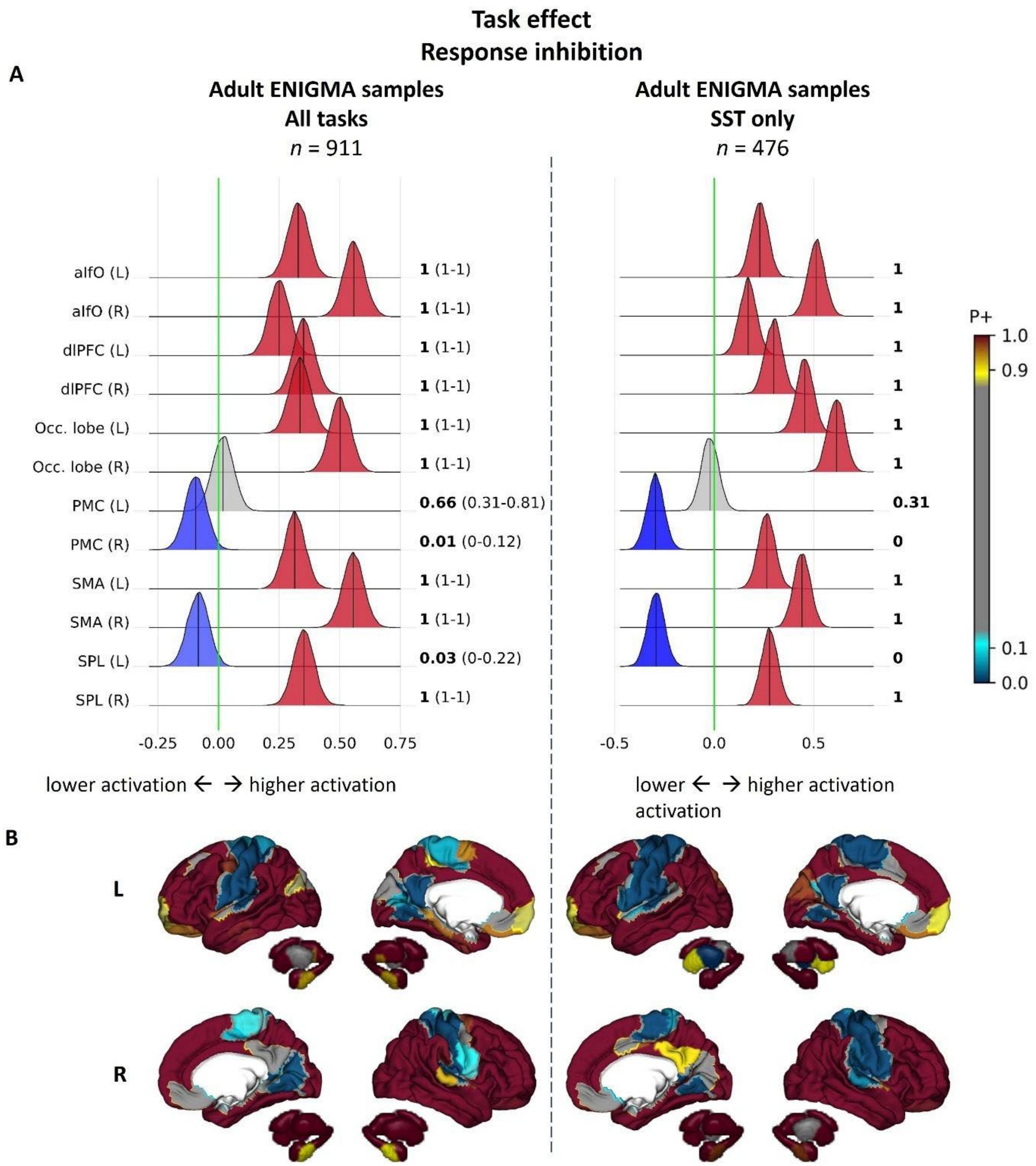

***Figure S6 - Comparison of main effect of response inhibition in adults from the ENIGMA-OCD sample across all tasks and in stop-signal tasks only*** (A) Region-of-interest results of group-level response inhibition contrast across individuals with OCD and HCs in all tasks (N tasks = 14, n participants = 911), and in only stop-signal tasks (N tasks = 10, n participants = 476). (B) Whole-brain analyses of response inhibition. aI/fO = anterior insula/frontal operculum; dlPFC = dorsolateral prefrontal cortex; L = left; Occ = occipital; PMC = primary motor cortex; R = right; SMA = supplementary motor area; SPL = superior parietal lobule; SST = stop-signal task.

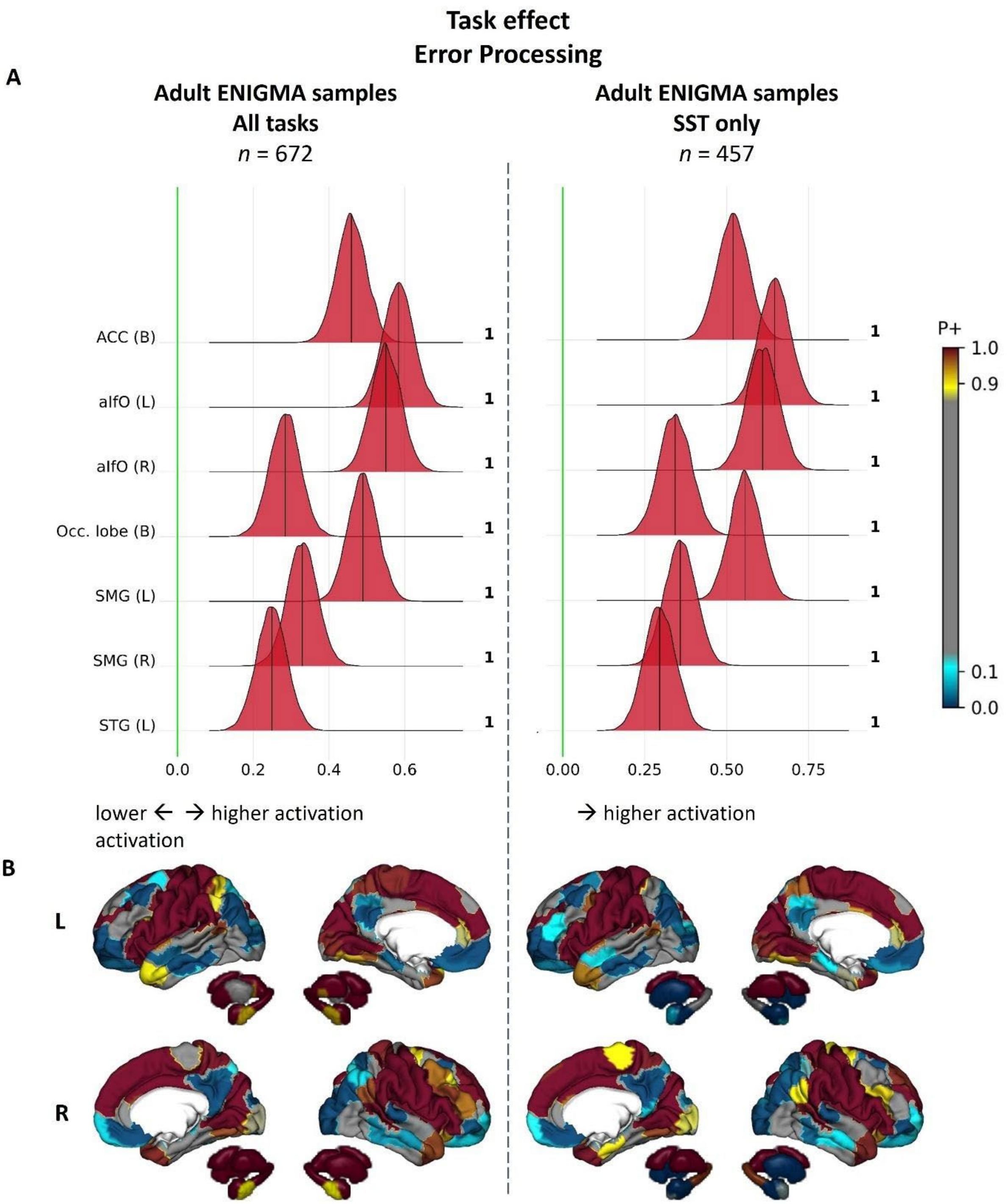

***Figure S7 - Comparison of main effect of error processing in adults from the ENIGMA-OCD sample across all tasks and in stop-signal tasks only*** (A) Region-of-interest results of group-level error processing contrast across individuals with OCD and HCs in all tasks (N tasks = 12, n participants = 672), and in only stop-signal tasks (N tasks = 10, n participants = 457). (B) Whole-brain analyses of error processing. ACC = anterior cingulate cortex; aI/fO = anterior insula/frontal operculum; L = left; Occ = occipital; R = right; SMG = supramarginal gyrus; STG = superior temporal gyrus; SST = stop-signal task.

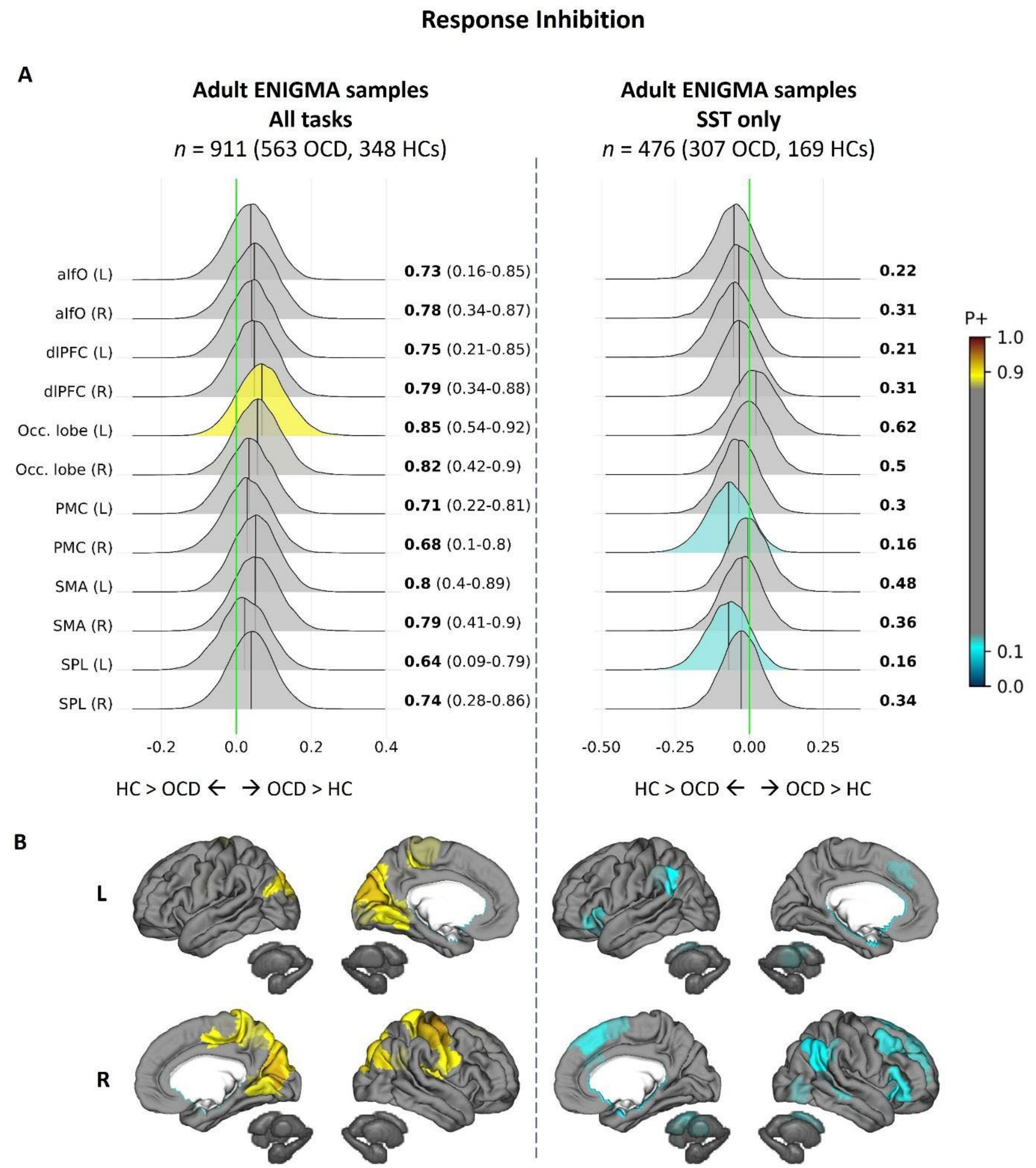

***Figure S8 - Comparison of case-control differences in response inhibition in adults from the ENIGMA-OCD sample across all tasks and in stop-signal tasks only*** (A) Region-of-interest results of response inhibition differences between individuals with OCD and HCs in all tasks (N tasks = 14, n participants = 911), and in only stop-signal tasks (N tasks = 10, n participants = 476). (B) Whole-brain analyses of response inhibition. aI/fO = anterior insula/frontal operculum; dlPFC = dorsolateral prefrontal cortex; L = left;  Occ = occipital; PMC = primary motor cortex; R = right; SMA = supplementary motor area; SPL = superior parietal lobule; SST = stop-signal task.

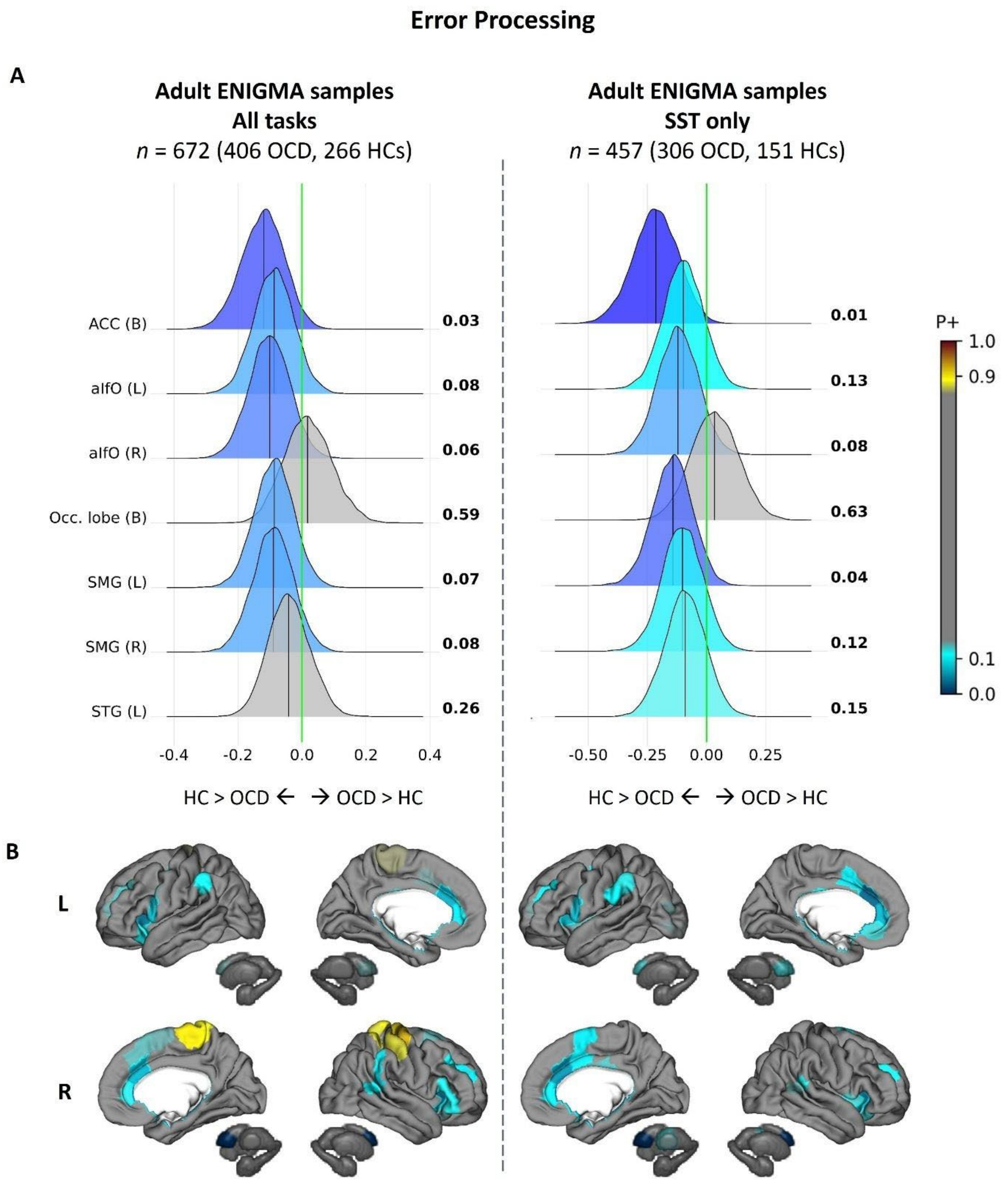

***Figure S9 - Comparison of case-control differences in error processing in adults from the ENIGMA-OCD sample across all tasks and in stop-signal tasks only*** (A) Region-of-interest results of error processing differences between individuals with OCD and HCs in all tasks (N tasks = 12, n participants = 672), and in only stop-signal tasks (N tasks = 10, n participants = 457). (B) Whole-brain analyses of error processing. ACC = anterior cingulate cortex; aI/fO = anterior insula/frontal operculum; L = left; Occ = occipital; R = right; SMG = supramarginal gyrus; STG = superior temporal gyrus; SST = stop-signal task.

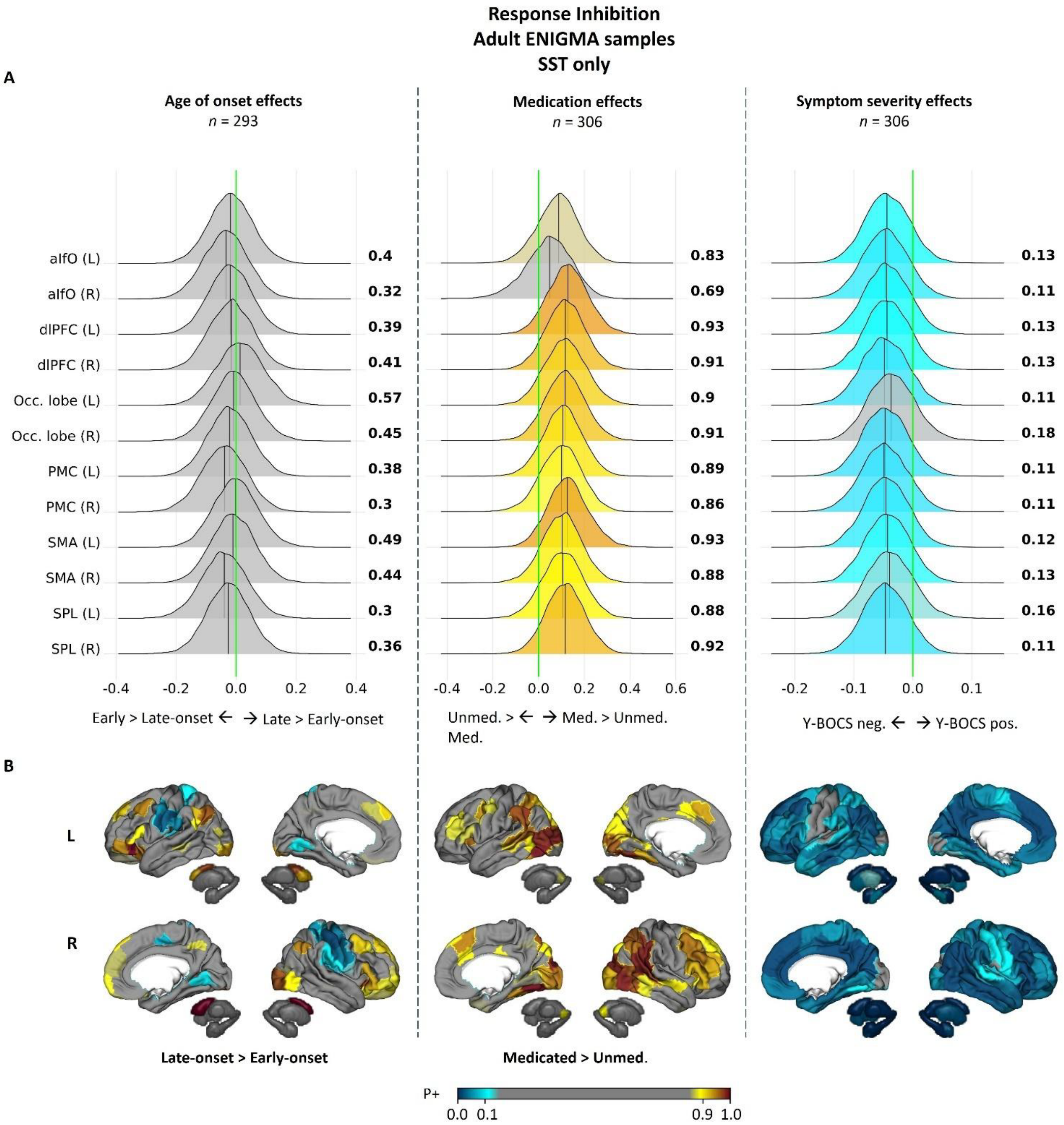

***Figure S10 - Effects of clinical features of OCD on response inhibition in adults from the ENIGMA-OCD sample selectively in stop-signal tasks*** (A) Region-of-interest effects of age of onset, medication status, and symptom severity on activation during response inhibition. (B) Whole-brain analyses of response inhibition. aI/fO = anterior insula/frontal operculum; dlPFC = dorsolateral prefrontal cortex; L = left; Occ = occipital; PMC = primary motor cortex; R = right; SMA = supplementary motor area; SPL = superior parietal lobule; SST = stop-signal task; Y-BOCS = Yale-Brown Obsessive-Compulsive Scale.

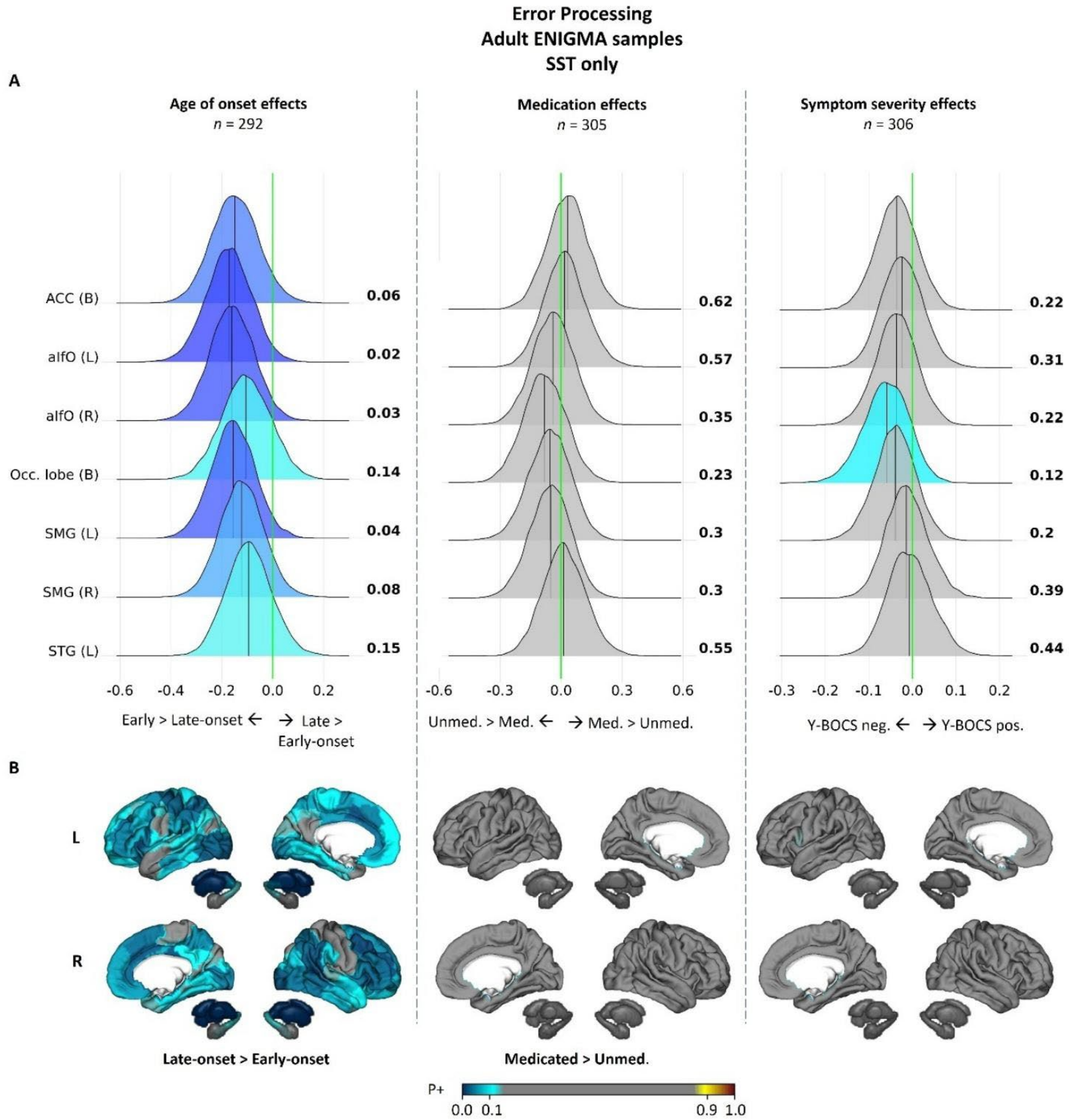

***Figure S11 - Effects of clinical features of OCD on error processing in adults from the ENIGMA-OCD sample selectively in stop-signal tasks*** (A) Region-of-interest effects of age of onset, medication status, and symptom severity on activation during error processing. (B) Whole-brain analyses of error processing. ACC = anterior cingulate cortex; aI/fO = anterior insula/frontal operculum; L = left; Occ = occipital; R = right; SMG = supramarginal gyrus; STG = superior temporal gyrus; SST = stop-signal task; Y-BOCS = Yale-Brown Obsessive-Compulsive Scale.

#### *Figure S12 - Whole-brain Schaefer 200-parcel cortical atlas & Melbourne 32 subcortical atlas Bayesian multilevel results*

The full output of the whole-brain Bayesian multilevel models presented in Figures 1-3 of the manuscript and Figures 1B-11B of this supplement can be retrieved from the online repository doi.org/10.5281/zenodo.17141947.

For each of the two contrasts of interest: 1) Response inhibition and 2) Error processing, the following models are obtained: Intercept model; Diagnosis group effect; Age of onset effect – Late-onset vs. HC – Early-onset vs. HC – Late-onset vs. Early-onset; Medication effect – Unmedicated vs. HC – Medicated vs. HC – Medicated vs. Unmedicated; OCD severity effect.

***Figure S13 - Sensitivity leave-one-sample-out analyses for Bayesian multilevel region-of-interest results for response inhibition contrast in adult ENIGMA-OCD samples***

*Group effect*

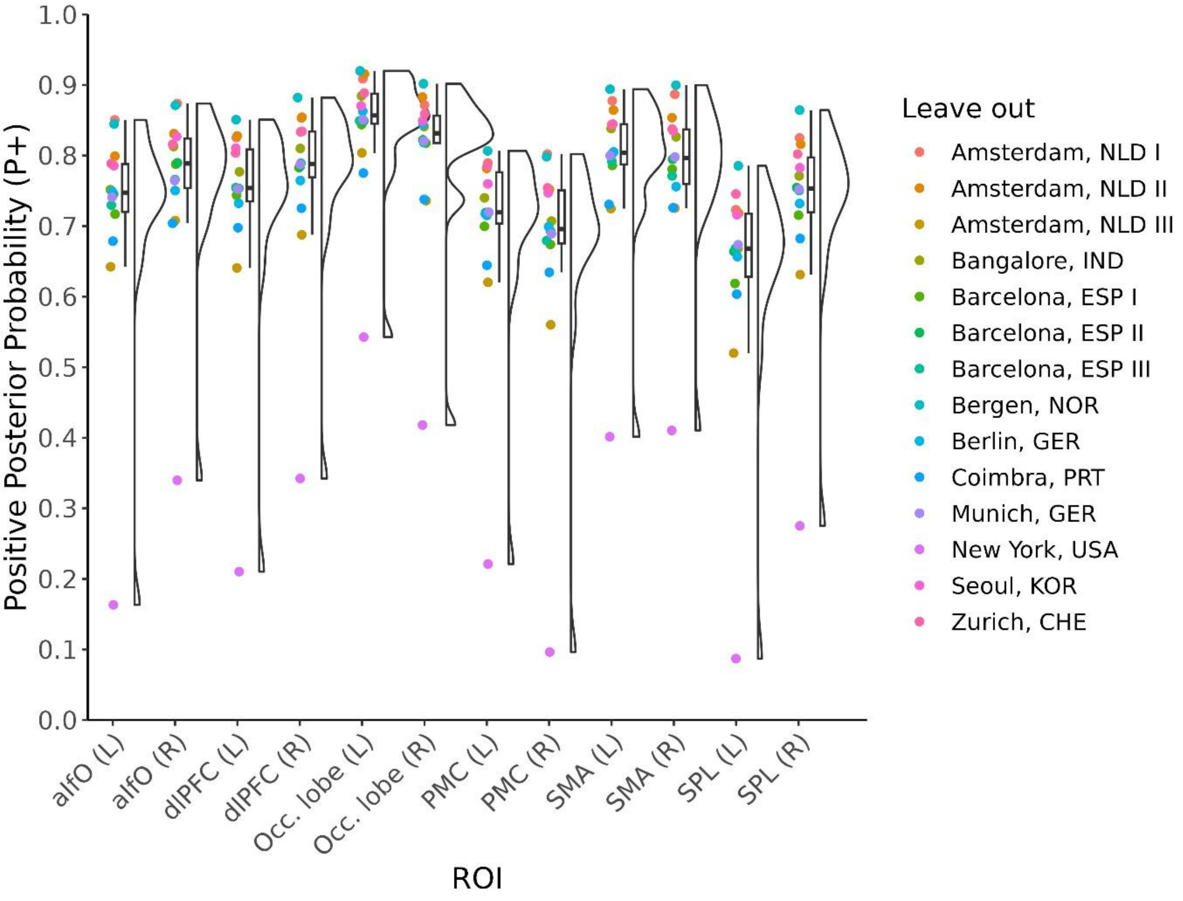

*Age of onset effect – Late-onset vs. HCs*

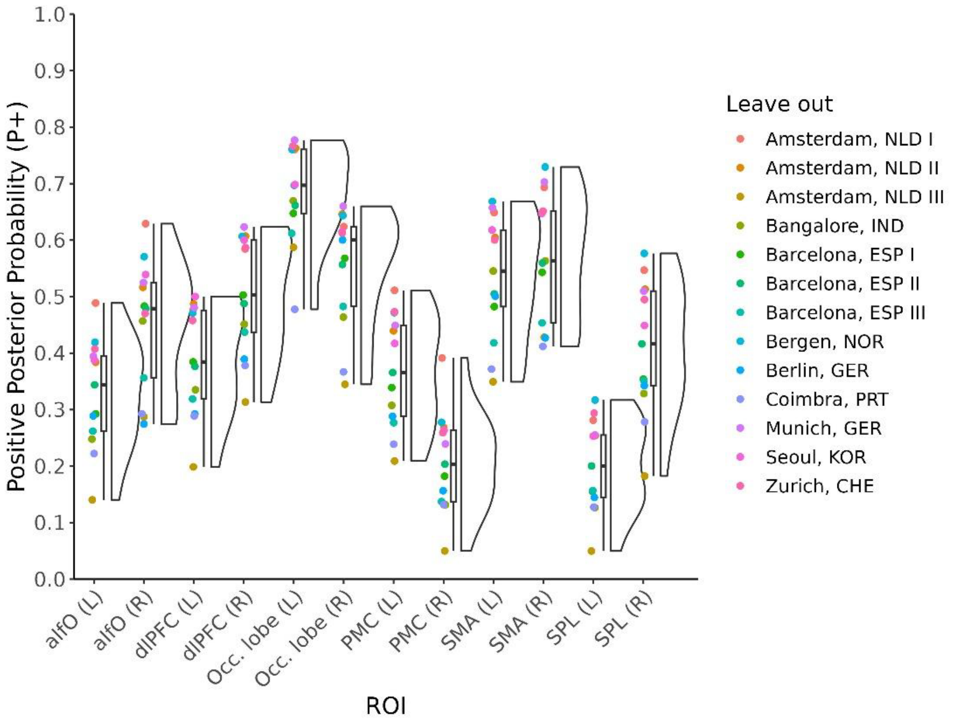

*Age of onset effect – Early-onset vs. HCs*

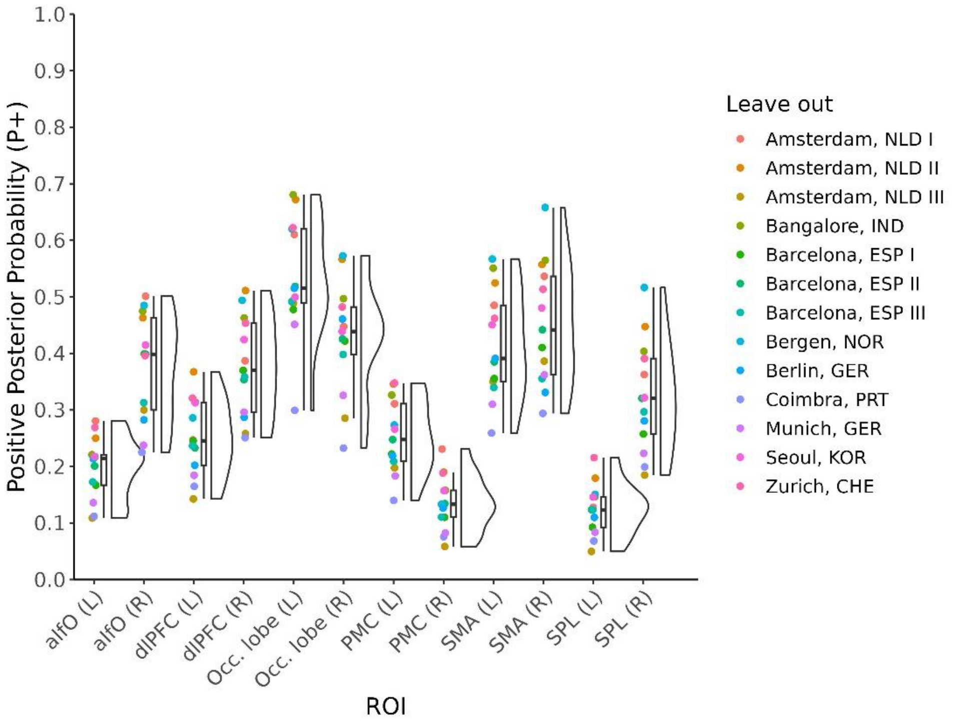

*Age of onset effect – Late-onset vs. Early-onset*

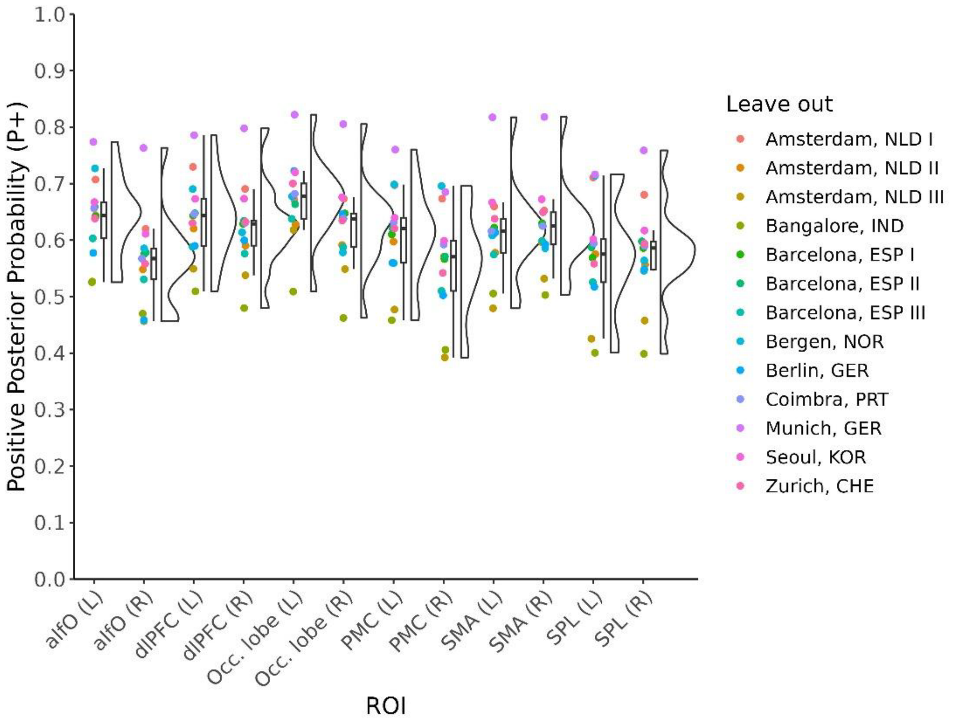

*Medication effect – Unmedicated vs. HCs*

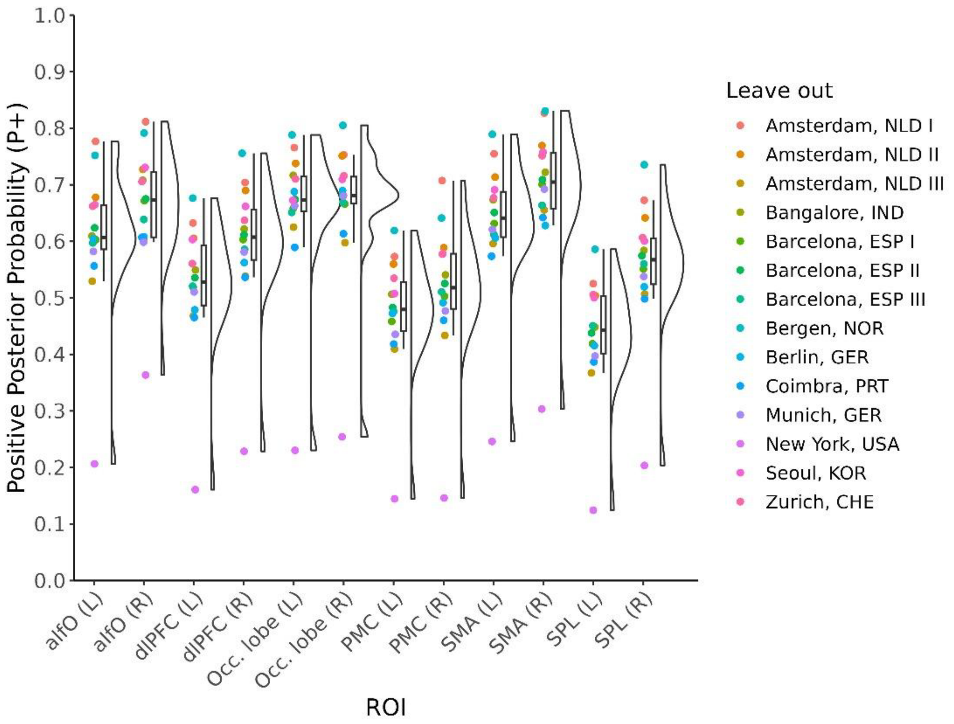

*Medication effect – Medicated vs. HCs*

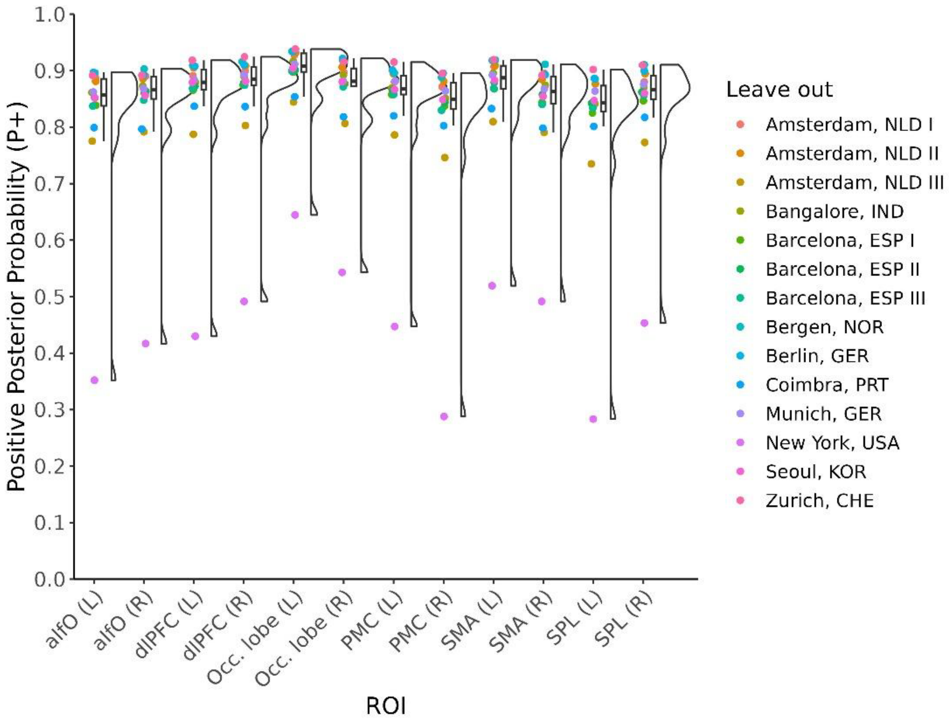

*Medication effect – Medicated vs. Unmedicated*

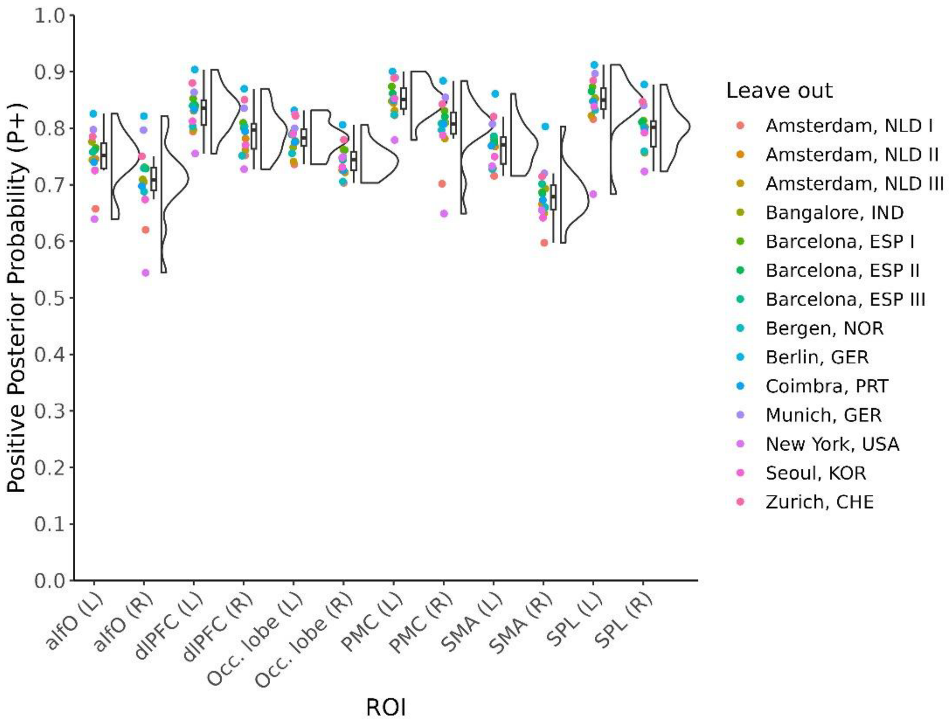

*Severity effect*

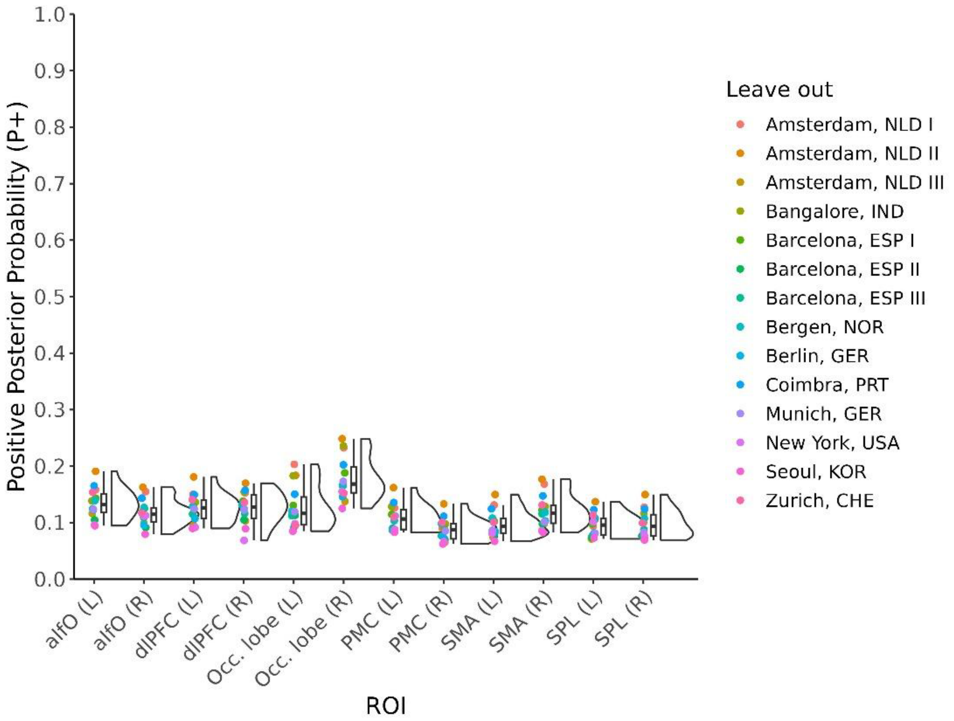

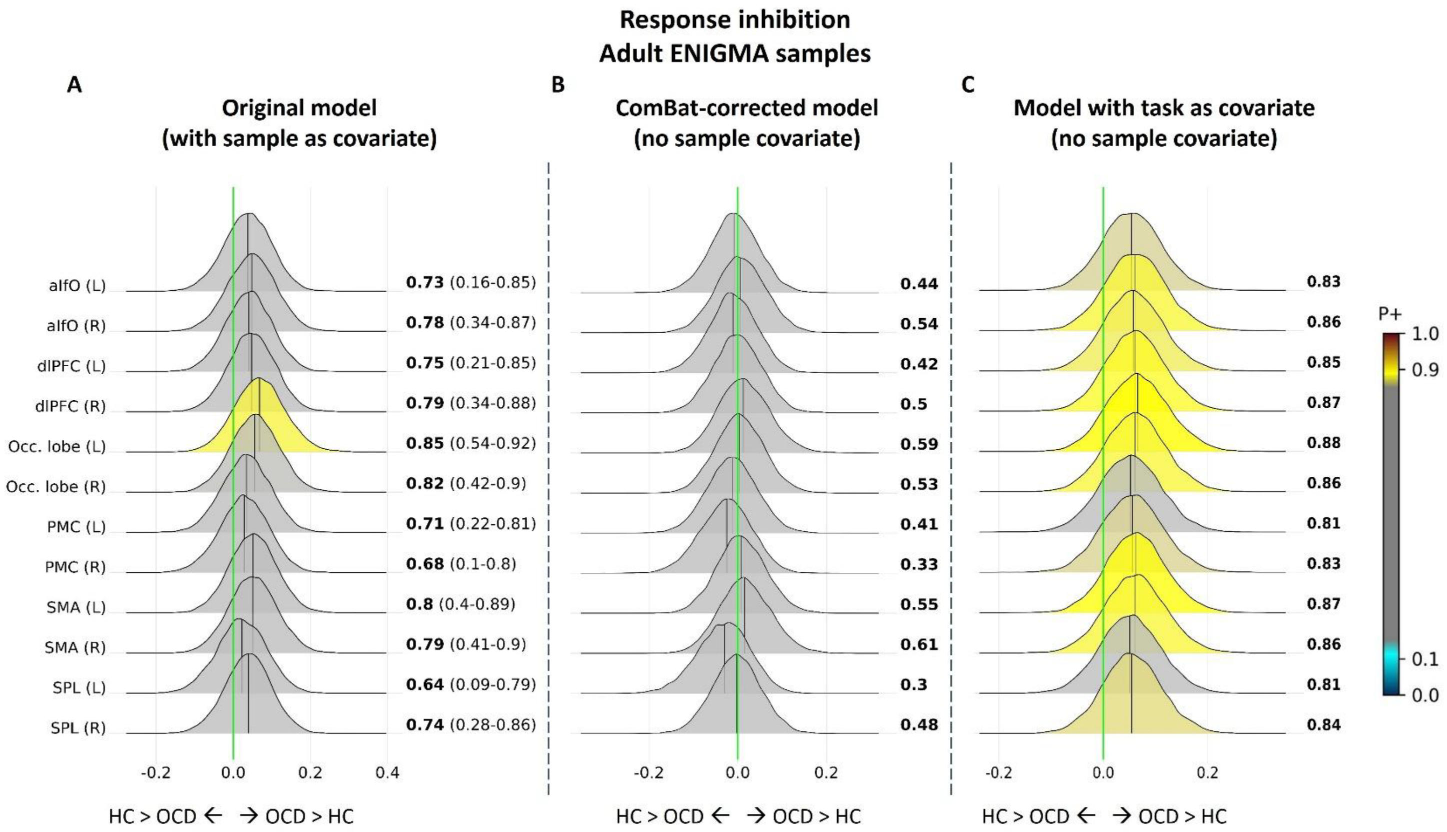

***Figure S14 - Sensitivity analyses of results across different sample and task correction methods*** (A) Original region-of-interest results from Bayesian multilevel analyses of case-control effects in response inhibition contrast. (B) Results after ComBat harmonization to correct for sample (batch) effects, with diagnosis, age, and sex included as biological covariates in the harmonization; the Bayesian model was run on the corrected data without including sample as a covariate. (C) Results from the original dataset with task type included as a covariate in the Bayesian model instead of sample to account for task-related variance. ACC = anterior cingulate cortex; aI/fO = anterior insula/frontal operculum; dlPFC = dorsolateral prefrontal cortex; L = left; Occ = occipital; PMC = primary motor cortex; R = right; SMA = supplementary motor area; SMG = supramarginal gyrus; SPL = superior parietal lobule; STG = superior temporal gyrus.

**Frequentist statistics**

Using a frequentist multilevel mixed effects model with random intercepts for sample and subject (instead of the Bayesian multilevel model used in the main analysis), we estimated response inhibition-related differences in activation between individuals with OCD and HCs (Table S4). Children with OCD from the ENIGMA-OCD sample showed significantly weaker activation in the right dlPFC compared to HCs (Z = -2.071, p = 0.039), while those from the ABCD sample showed significantly weaker activation in the right aI/fO (Z = -2.075, p = 0.038). During error processing, adults with OCD had significantly weaker activation in the right SMG relative to HCs (Table S5). No other effects were found to be significant.

**Table S5 – Results of frequentist multilevel analysis of activation differences in predefined ROIs between individuals with OCD and HC during response inhibition**
ACC = anterior cingulate cortex; aI/fO = anterior insula/frontal operculum; dlPFC = dorsolateral prefrontal cortex; Occ = occipital; PMC = primary motor cortex; SMA = supplementary motor area; SMG = supramarginal gyrus; SPL = superior parietal lobule; STG = superior temporal gyrus.

| **Region** | **B [SE]** | ***Z*** | **Cohen's *d*** | ***p* value** |
|---|---|---|---|---|
| **Adult ENIGMA-OCD samples** | | | | |
| aIfO (L) | 0.031 [0.081] | 0.382 | 0.035 | 0.703 |
| aIfO (R) | 0.138 [0.081] | 1.700 | 0.154 | 0.089 |
| dlPFC (L) | 0.067 [0.081] | 0.824 | 0.075 | 0.410 |
| dlPFC (R) | 0.109 [0.081] | 1.341 | 0.122 | 0.180 |
| Occ. lobe (L) | 0.068 [0.081] | 0.836 | 0.076 | 0.403 |
| Occ. lobe (R) | 0.025 [0.081] | 0.308 | 0.028 | 0.758 |
| PMC (L) | -0.082 [0.081] | -1.007 | -0.092 | 0.314 |
| PMC (R) | 0.005 [0.081] | 0.063 | 0.006 | 0.950 |
| SMA (L) | 0.092 [0.081] | 1.135 | 0.103 | 0.256 |
| SMA (R) | 0.137 [0.081] | 1.691 | 0.154 | 0.091 |
| SPL (L) | -0.043 [0.081] | -0.526 | -0.048 | 0.599 |
| SPL (R) | 0.027 [0.081] | 0.329 | 0.030 | 0.742 |
| **Pediatric ENIGMA-OCD samples** | | | | |
| aIfO (L) | -0.137 [0.118] | -1.160 | -0.210 | 0.246 |
| aIfO (R) | -0.151 [0.118] | -1.278 | -0.231 | 0.202 |
| dlPFC (L) | -0.083 [0.118] | -0.704 | -0.128 | 0.481 |
| dlPFC (R) | -0.245 [0.118] | -2.071 | -0.375 | 0.039 * |
| Occ. lobe (L) | 0.052 [0.118] | 0.441 | 0.080 | 0.659 |
| Occ. lobe (R) | 0.2 [0.118] | 1.691 | 0.306 | 0.091 |
| PMC (L) | 0.011 [0.118] | 0.090 | 0.016 | 0.929 |
| PMC (R) | 0.056 [0.118] | 0.470 | 0.085 | 0.638 |
| SMA (L) | -0.188 [0.118] | -1.590 | -0.288 | 0.112 |
| SMA (R) | -0.162 [0.118] | -1.373 | -0.248 | 0.170 |
| SPL (L) | 0.043 [0.118] | 0.366 | 0.066 | 0.715 |
| SPL (R) | 0.048 [0.118] | 0.404 | 0.073 | 0.686 |

| Pediatric ABCD samples | | | | |
|---|---|---|---|---|
| aIfO (L) | -0.053 [0.057] | -0.929 | -0.093 | 0.353 |
| aIfO (R) | -0.119 [0.057] | -2.075 | -0.207 | 0.038 * |
| dlPFC (L) | 0.035 [0.057] | 0.601 | 0.060 | 0.548 |
| dlPFC (R) | -0.076 [0.057] | -1.316 | -0.131 | 0.188 |
| Occ. lobe (L) | -0.026 [0.057] | -0.446 | -0.044 | 0.655 |
| Occ. lobe (R) | 0.018 [0.057] | 0.315 | 0.031 | 0.753 |
| PMC (L) | 0.039 [0.057] | 0.673 | 0.067 | 0.501 |
| PMC (R) | 0.008 [0.057] | 0.140 | 0.014 | 0.888 |
| SMA (L) | -0.034 [0.057] | -0.583 | -0.058 | 0.560 |
| SMA (R) | -0.107 [0.057] | -1.865 | -0.186 | 0.062 |
| SPL (L) | -0.035 [0.057] | -0.605 | -0.060 | 0.545 |
| SPL (R) | -0.089 [0.057] | -1.554 | -0.155 | 0.120 |

* = statistically significant at an alpha level of 0.05

**Table S6 – Results of frequentist multilevel analysis of activation differences in predefined ROIs between individuals with OCD and HC during error processing**
ACC = anterior cingulate cortex; aI/fO = anterior insula/frontal operculum; dlPFC = dorsolateral prefrontal cortex; Occ = occipital; PMC = primary motor cortex; SMA = supplementary motor area; SMG = supramarginal gyrus; SPL = superior parietal lobule; STG = superior temporal gyrus.

| Region | B [SE] | *Z* | Cohen's *d* | *p* value |
|---|---|---|---|---|
| **Adult ENIGMA-OCD samples** | | | | |
| ACC (B) | -0.127 [0.071] | -1.792 | -0.187 | 0.073 |
| aIfO (L) | -0.074 [0.071] | -1.046 | -0.109 | 0.295 |
| aIfO (R) | -0.111 [0.071] | -1.574 | -0.164 | 0.115 |
| Occ. lobe (B) | 0.075 [0.071] | 1.055 | 0.110 | 0.291 |
| SMG (L) | -0.082 [0.071] | -1.160 | -0.121 | 0.246 |
| SMG (R) | -0.142 [0.071] | -2.011 | -0.210 | 0.044 * |
| STG (L) | -0.012 [0.071] | -0.173 | -0.018 | 0.863 |
| **Pediatric ENIGMA-OCD samples** | | | | |
| ACC (B) | 0.117 [0.128] | 0.914 | 0.178 | 0.361 |
| aIfO (L) | 0.144 [0.128] | 1.126 | 0.219 | 0.261 |
| aIfO (R) | 0.161 [0.128] | 1.253 | 0.244 | 0.211 |
| Occ. lobe (B) | 0.01 [0.128] | 0.078 | 0.015 | 0.938 |
| SMG (L) | 0.028 [0.128] | 0.218 | 0.042 | 0.828 |
| SMG (R) | 0.1 [0.128] | 0.778 | 0.152 | 0.437 |
| STG (L) | -0.047 [0.128] | -0.365 | -0.071 | 0.715 |
| **Pediatric ABCD samples** | | | | |
| ACC (B) | -0.013 [0.058] | -0.230 | -0.023 | 0.818 |
| aIfO (L) | 0.056 [0.058] | 0.967 | 0.095 | 0.333 |
| aIfO (R) | 0.091 [0.058] | 1.577 | 0.155 | 0.115 |
| Occ. lobe (B) | -0.051 [0.058] | -0.874 | -0.086 | 0.382 |
| SMG (L) | 0.081 [0.058] | 1.399 | 0.138 | 0.162 |
| SMG (R) | 0.047 [0.058] | 0.811 | 0.080 | 0.417 |
| STG (L) | 0.001 [0.058] | 0.015 | 0.001 | 0.988 |

* = statistically significant at an alpha level of 0.05

### References

1. de Wit SJ, de Vries FE, van der Werf YD, Cath DC, Heslenfeld DJ, Veltman EM, et al. Presupplementary motor area hyperactivity during response inhibition: a candidate endophenotype of obsessive-compulsive disorder. Am J Psychiatry. 2012;169(10):1100-8.
2. Fitzsimmons SMDD, Postma T, van Campen AD, Vriend C, Batelaan NM, van Oppen P, et al. TMS-induced plasticity improving cognitive control in OCD I: Clinical and neuroimaging outcomes from a randomised trial of rTMS for OCD. medRxiv. 2023:2023.11.04.23298100.
3. van der Straten A, Bruin W, van de Mortel L, ten Doesschate F, Merkx MJM, de Koning P, et al. Pharmacological and Psychological Treatment Have Common and Specific Effects on Brain Activity in Obsessive-Compulsive Disorder. Depression and Anxiety. 2024;2024(1):6687657.
4. Huyser C, Veltman DJ, Wolters LH, de Haan E, Boer F. Developmental aspects of error and high-conflict-related brain activity in pediatric obsessive–compulsive disorder: a fMRI study with a Flanker task before and after CBT. Journal of Child Psychology and Psychiatry. 2011;52(12):1251-60.
5. Suñol M, Martínez-Zalacaín I, Picó-Pérez M, López-Solà C, Real E, Fullana MÀ, et al. Differential patterns of brain activation between hoarding disorder and obsessive-compulsive disorder during executive performance. Psychological Medicine. 2020;50(4):666-73.
6. Grützmann R, Kaufmann C, Wudarczyk OA, Balzus L, Klawohn J, Riesel A, et al. Error-Related Brain Activity in Patients With Obsessive-Compulsive Disorder and Unaffected First-Degree Relatives: Evidence for Protective Patterns. Biological Psychiatry Global Open Science. 2022;2(1):79-87.
7. Stern ER, Brown C, Ludlow M, Shahab R, Collins K, Lieval A, et al. The buildup of an urge in obsessive–compulsive disorder: Behavioral and neuroimaging correlates. Human Brain Mapping. 2020;41(6):1611-25.
8. Hollestein V, Buitelaar JK, Brandeis D, Banaschewski T, Kaiser A, Hohmann S, et al. Developmental changes in fronto-striatal glutamate and their association with functioning during inhibitory control in autism spectrum disorder and obsessive compulsive disorder. NeuroImage: Clinical. 2021;30:102622.
9. Kang D-H, Jang JH, Han JY, Kim J-H, Jung WH, Choi J-S, et al. Neural correlates of altered response inhibition and dysfunctional connectivity at rest in obsessive–compulsive disorder. Progress in Neuro-Psychopharmacology and Biological Psychiatry. 2013;40:340-6.
10. Iannaccone R, Hauser TU, Staempfli P, Walitza S, Brandeis D, Brem S. Conflict monitoring and error processing: new insights from simultaneous EEG-fMRI. Neuroimage. 2015;105:395-407.
11. Norman LJ, Taylor SF, Liu Y, Radua J, Chye Y, De Wit SJ, et al. Error Processing and Inhibitory Control in Obsessive-Compulsive Disorder: A Meta-analysis Using Statistical Parametric Maps. Biol Psychiatry. 2019;85(9):713-25.

12. Chen G, Xiao Y, Taylor PA, Rajendra JK, Riggins T, Geng F, et al. Handling Multiplicity in Neuroimaging Through Bayesian Lenses with Multilevel Modeling. Neuroinformatics. 2019;17(4):515-45.
